# Sequential predictive learning is a unifying theory for hippocampal representation and replay

**DOI:** 10.1101/2024.04.28.591528

**Authors:** Daniel Levenstein, Aleksei Efremov, Adel Halawa, Roy Henha Eyono, Jonathan Cornford, Adrien Peyrache, Blake Richards

## Abstract

The hippocampus represents an animal’s position in the environment and generates “replay” proposed to support recall, planning, and memory formation. Recently, it’s been found that artificial neural networks trained to predict sensory inputs develop spatially-tuned cells, however, whether predictive learning can account for the ability to produce replay, or the hippocampus’ representation of non-spatial variables, is unknown. Here, we find that offline replay only emerged in recurrent neural networks that used head-direction information to predict multi-step observation sequences, which promoted the formation of a continuous attractor reflecting the geometry of the environment. These offline trajectories were able to autonomously replay recently experienced locations, generate novel paths, and could be directed by a virtual head direction signal. Further, these networks developed sweeping representations of future positions and represented non-spatial task-relevant variables. These results demonstrate how hippocampal-like operations can emerge in neural networks engaged in sequential predictive learning, providing a unifying theory for hippocampal functions.

## Main

The mammalian hippocampus has been implicated in seemingly disparate cognitive processes, including navigation ^1–4^, memory ^5–8^, planning ^9–11^, and imagination ^12,13^. This functional diversity appears to rely on the interplay between two distinct modes of operation ^14^. The first is an input-driven “online” mode during active behavior in the environment ^15^. In the online mode, neural activity shows prominent theta rhythms ^15^, cells are tuned to an animal’s position ^1^ and non-spatial task features ^16–20^, and population activity lies on a low-dimensional manifold that reflects environmental and task structure ^21–24^ (**Figure 1a**). The second is an “offline” mode during periods of behavioral quiescence and sleep ^15^. In the offline mode, irregular neural activity “replays’’ realistic trajectories through the environment ^25–27^, including sequences of positions from previous experiences ^26^ and generative trajectories of untaken paths ^28–30^. Importantly, replay is internally generated in the hippocampus ^31–35^, but may be influenced by its inputs ^32,35,36^ such as the head direction system which conveys a coherent but randomly drifting signal during sleep ^37^ (**Figure 1b**). In downstream structures, replay engages sensory representations that were active when the animal visited that position ^38^. While multiple theories have been proposed to account for the hippocampus’ computational capacities^39–41^, a unified account of these modes of operation is still lacking. In particular, we lack a model for how the hippocampus can learn a distributed representation of the environment from sensory information during the online mode, and generate offline activity that meets three key desiderata: (1) A coherent representation of spatial position in the absence of sensory input ^42^; (2) Autonomous generation of sequences of positions that produce plausible trajectories through the environment ^26,29^; and (3) Output of associated sensory inputs from learned associations at those positions ^38^ (**Figure 1c**).

**Figure 1:**
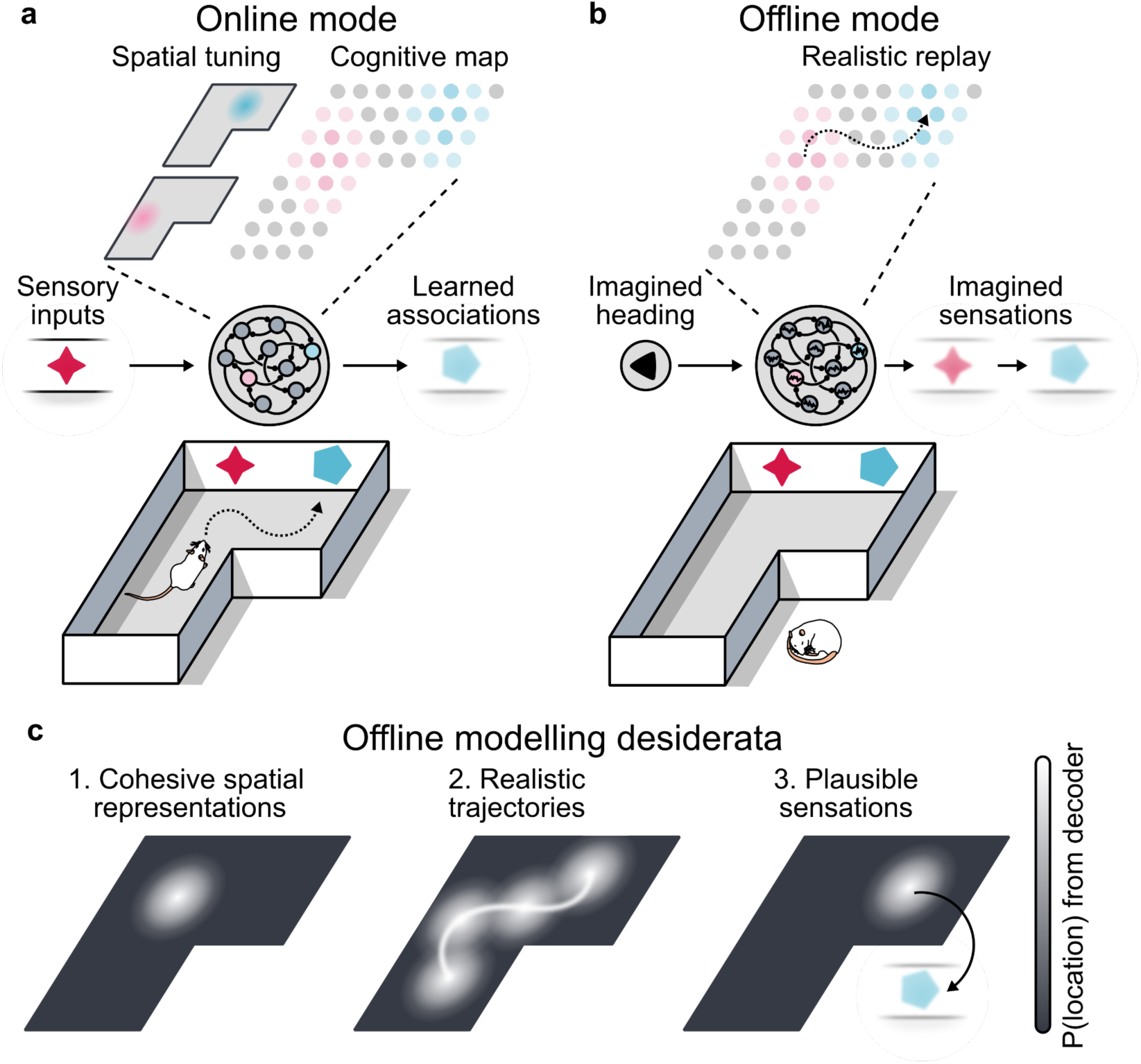
Hippocampal representation and replay. **a:** During active behavior in the environment, the hippocampus is in an online mode of activity. Based on sensory inputs, it contains spatially tuned cells (e.g. two cells active in red and blue areas, respectively) and a cognitive map representation of the environment, which it uses to simulate the immediate consequences of future behavior and output learned associations to the rest of the brain. **b:** During consummatory and quiescent behavior, the hippocampus is in an offline mode of activity. It generates simulated “replay” trajectories, which project learned associations to the rest of the brain, possibly under the influence of an imagined head direction signal. **c:** Replay trajectories are sequences of internally-generated coherent representations of spatial locations, which follow realistic trajectories in the environment, and output learned associations of those locations to the rest of the brain.

A classic theory to explain hippocampal representation and replay is that the hippocampal CA3 subregion implements a continuous attractor neural network (CANN, ^41,43,44^). CANNs represent space with an attractive neural manifold ^45^, in which recurrent connectivity maintains a localized bump of activity even in the absence of input. Indeed, CANN models can account for a number of experimental observations during both online and offline activity. These include: 1) the presence of spatially-tuned cells ^1^, 2) population activity that lies on a low-dimensional manifold ^21–23^, and 3) internally-generated offline trajectories that can range from diffusive motion ^46^ to memorized trajectories ^47^. However, traditional CANN models rely on the pre-assignment of neurons to spatial locations or hand-tuned recurrent connectivity ^43,44,48^, and models that learn CANNs rely on signals with pre-existing tuning for allocentric space ^49–51^. Thus, while CANN models can account for spatial representation and replay, they do not account for the hippocampus’ non-spatial representations or how these properties can emerge in a network, i.e. how they can be learned purely from egocentric sensory inputs, such as images, and actions. Further, CANNs cannot account for two other facets of hippocampal function. The first is the association of viewpoint-dependent sensory information with a viewpoint-independent representation of space. The second is the predominance of weakly tuned cells which are critical for the low dimensional structure of population activity in the hippocampus ^23,52^ and its distributed representation of space ^53,54^.

One promising alternative is predictive learning. Where traditional CANN models rely on preexisting knowledge of space, a predictive learning network uses the difference between predictions and inputs to modify connection strengths, and thus its representational capacities emerge as a consequence of learning to predict. Computational models have demonstrated that learning to predict upcoming sensory inputs leads to the emergence of spatially-tuned units in artificial neural networks ^55,56^. This spatial tuning, when analyzed at the population level, can exhibit low-dimensional manifolds that reflect the environment, similar to the hippocampus ^21–24,55,56^. In addition, hippocampal place fields resemble predictive representations used in reinforcement learning ^39,57^, and the hippocampus has been shown to encode expectations about upcoming stimuli ^58,59^ and prediction errors ^60,61^. Indeed, alternative theories have proposed that the hippocampus is a predictive or sequence generation network ^40,62–65^, based on its anatomy ^62,63^, recurrent circuit connectivity of CA3 ^66,67^, and observations that neuronal activity does not just encode an animals current location, but rather, “sweeps’’ in a manner that represents trajectories of possible future positions ^68–70^. However, spatially-tuned units are not unique to predictive learning. Grid cells and toroidal manifolds emerge in networks trained on path integration or spatial classification tasks ^49,50,71,72^, though these are models of medial entorhinal cortex that require spatial position information in their training objective. Thus, it remains unknown if a single learning objective, applied to a distributed recurrent network receiving only egocentric sensory inputs, can account for the hippocampus’ ability to generate offline replay or to represent non-spatial, task-relevant variables.

To determine the potential for predictive learning to unify the different aspects of hippocampal function, we trained recurrent neural networks (RNNs) to predict egocentric sensory inputs as an agent moved through a simulated environment (**Figure 2a**, Methods). This allowed us to compare the representations learned by different predictive models (**Figure S1**), in a setting in which agents received spatially ambiguous sensory input. It also allowed us to study the models’ offline generation capabilities in the absence of sensory input. We found that an attractor-based cognitive map capable of generating offline replay can emerge as a consequence of predictive learning, but only when networks use recurrent connections to predict multi-step sequences of sensory input using an orienting (head direction) signal. The resulting networks contained a distributed representation of the agent’s environment with a diverse population of spatially-tuned cells, produced a coherent representation of spatial trajectories in the absence of sensory input that did not rely on a sensory loop from predictions and could be influenced by recent experience or fictitious actions, and represented task-relevant non-spatial variables as well as the agent’s position. In sum, sequential predictive learning can account for both online representations and offline replay in the hippocampus, and is a candidate theory to unify three prominent views of the hippocampus: 1) the hippocampus is a predictive map ^39^, 2) the hippocampus is a CANN ^41,43,44,73^, and 3) the hippocampus is a sequence generator ^40^.

**Figure 2:**
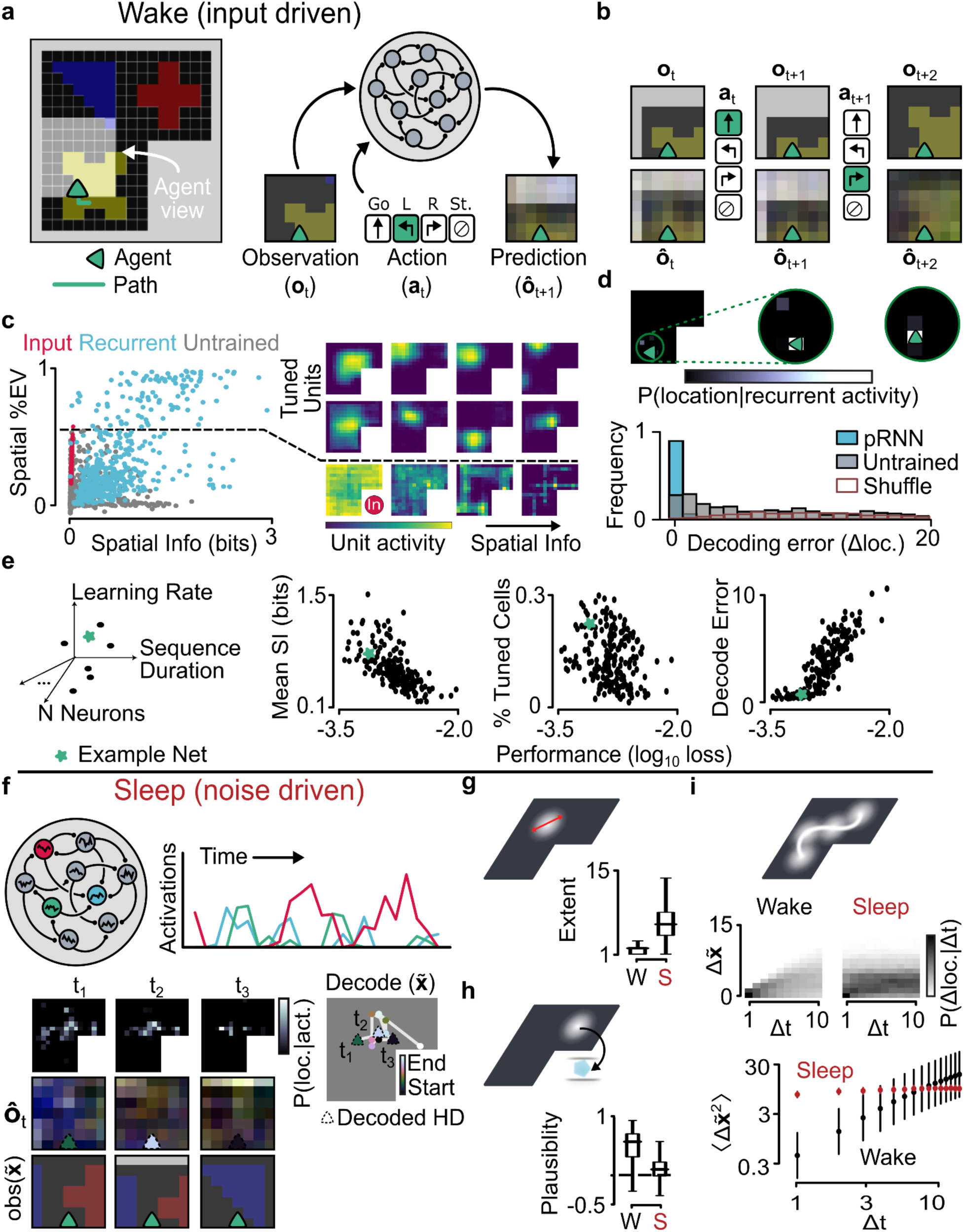
Next-step predictive learning produces spatially tuned cells but not replay. **a:** (Left) The agent moves around the gridworld environment with a visual field corresponding to the 7×7 grid of tiles in front of it. Gray tiles represent impassable walls, while colored floor tiles provide visual cues. (Right) a predictive RNN (pRNN) is trained to use the current observation and action to predict the observation in the next timestep. **b:** Example sequence of observations and predicted observations from a trained network. **c:** (Left) Explained variance by spatial position and spatial information of all units in the example recurrent network, input units, and units from an untrained random network. (Right) spatial tuning curves from a selection of example tuned units (units with >50% variance explained by spatial position) with unimodal place fields, untuned units, and an example input unit. **d:** (Top) Decoded position from a linear decoder on hidden unit activity for the example three timesteps in B. (Bottom) distribution of spatial decoding error over a 1000-timestep trajectory in the environment, for the example pRNN and untrained network, compared to a position-shuffled control trajectory. **e:** Spatial representation (mean spatial information, percentage of tuned cells, linear decoding error) in a population of pRNNs with different hyperparameter values. Teal star reflects the example network in b-d. **f:** Sleep-like activity is generated in the network by removing the sensory input and driving each unit with noise. This activity is decoded using the linear decoder from d. Outputs from the network reflect the predicted “observations” associated with offline activity. **g**: The distribution of the decoded posterior’s spatial extent (a measure of how coherently the network is representing a single position) during wake and sleep (combined over 15 sleep epochs). Box indicates Q1, median, Q3; whiskers indicate data range. **h**: The distribution of plausibility of the predicted visual output from the network during wake and sleep. Plausibility is measured as the correlation between the visual output and true visual field at the decoded position and head direction. Box indicates Q1, median, Q3. Whiskers indicate data range. **i**: Spatial transition statistics of the decoded trajectories during sleep and wake. While the variance of agent’s position during wake increased with time (by construction), the decoded position during sleep incoherently shifted from location to location, irrespective of the position at the immediately preceding time step. Error bars indicate standard deviation over 15 sleep epochs.

### Next-step predictive learning produces spatially tuned cells but not replay

We simulated an agent in a “grid world” environment with visual cues consisting of different colors and patterns of floor tiles (**Figure 2a**). During a “wake” epoch, the agent took random actions that allowed for thorough exploration of the environment (Methods). At each location the agent received a 7×7 color image corresponding to the egocentric view of the floor and walls immediately in front of it (**Figure 2b**, **Supplemental Movie**). Critically, the inputs to this network were both spatially ambiguous and redundant (**Figure 2c, S2**) – individual sensory units had very little spatial tuning, multiple locations in the environment had the same sensory input, and the same position could have different sensory input depending on the agent’s head direction. The sequence of actions and visual inputs from each epoch were used to train a RNN to predict the next visual input at each timestep using backpropagation through time ^55^ (**Figure 2a, b, Figure S3a,** Methods).

After training, the network contained an allocentric representation of space (**Figure 2c,d**). The agent’s location was accurately decodable from the predictive RNN units via a linear classifier (**Figure 2b, d)**, Methods), a large part of the variance of the activity of many RNN units’ activity could be explained by the agent’s position (141±15/600 units with >50% variance explained by spatial position), and they developed spatial tuning curves that carried significantly more spatial information than was observed in the visual inputs or untrained networks (**Figure 2c**, Methods). Like cells in the hippocampus ^74^, units in the network had a skewed distribution of spatial information (**Figure S3b**), with a large number of weakly tuned cells and a small number of cells with strong/reliable spatial tuning.

The emergence of spatially tuned units was a robust property of predictive learning in a recurrent network. The trained RNNs had units with more spatial information than a random (untrained) RNN^75^ (**Figure 2c,d**) or identically trained feedforward networks (**Figure S4**), and the emergence of spatially tuned cells did not rely on the geometry of the environment (**Figure S5**), or the use of a discrete environment and action space (**Figure S6**). Because the emergent properties of neural network models can be sensitive to specific hyperparameter choices ^76^, we trained a population of 250 predictive networks, each with a different hyperparameter setting (seed, learning rate, sequence duration, back-propagation time-window, number of neurons, and neural time-scale; **Figure S7**). We found that the emergence of a spatial representation was strongly correlated with predictive performance across this population of RNNs (**Figure 2e**) and, critically, no network learned to predict well without developing spatially tuned cells. These results demonstrate that allocentric spatial tuning is a natural and robust consequence of learning to predict egocentric sensory information in recurrent neural networks.

We next tested if the predictive RNNs were able to generate plausible replay during a sleep-like state. Specifically, we removed the sensory and action inputs and increased the amount of noise above that used during training (uncorrelated Gaussian noise, see Methods), such that network dynamics was predominantly governed by recurrent connectivity (**Figure S8**). To identify the locations represented by the network during sleep, we used the decoder trained to decode position from the RNNs’ activity during wake (**Figure 2f**). We found that the network did not represent a spatially localized position (**Figure 2g**), the sensory “predictions” produced by offline activity did not correspond to the sensory input from the decoded viewpoint (i.e. position and head direction, **Figure 2h**), and that trajectories of represented locations did not conform to environmental statistics (**Figure 2I**). Instead, the trajectory frequently jumped around space and only visited a small number of disconnected locations in the environment, which were the same from trial to trial (**Figure S8**). This was in stark contrast to trajectories during wake, which were restricted to smooth transitions through the environment. The inability to produce replay was not a result of specific hyperparameter choices, as the networks were unable to produce reasonable trajectories regardless of the network hyperparameters or level of noise (**Figure S8**). Thus, learning to predict the next image robustly produced spatial tuning but did not produce the ability to intrinsically generate replay.

### A continuous attractor-based representation emerges from sequential predictive learning

In examining the population activity of the predictive RNNs, we observed that, despite the presence of spatially tuned units, activity in the RNN did not resemble a CANN (**Figure 3a,b**). That is, population activity during wake did not lie on a manifold that mirrored the spatial layout of the environment, and the activity during sleep converged to a single off-manifold fixed point. To quantify these observations, we developed two metrics: first, a spatial representational similarity analysis (sRSA) that measured the correlation between distance in neural space and distance in the environment (Methods); and second, a sleep-wake distance (S-W Dist) that measured the distance in neural space between activity during sleep and the wake manifold (Methods). We observed that the sRSA scores for the predictive RNNs were significantly lower than those for the CANNs (**Figure 3a,b**), while their S-W Dist was significantly higher, indicating that population activity lied outside the wake manifold.

**Figure 3:**
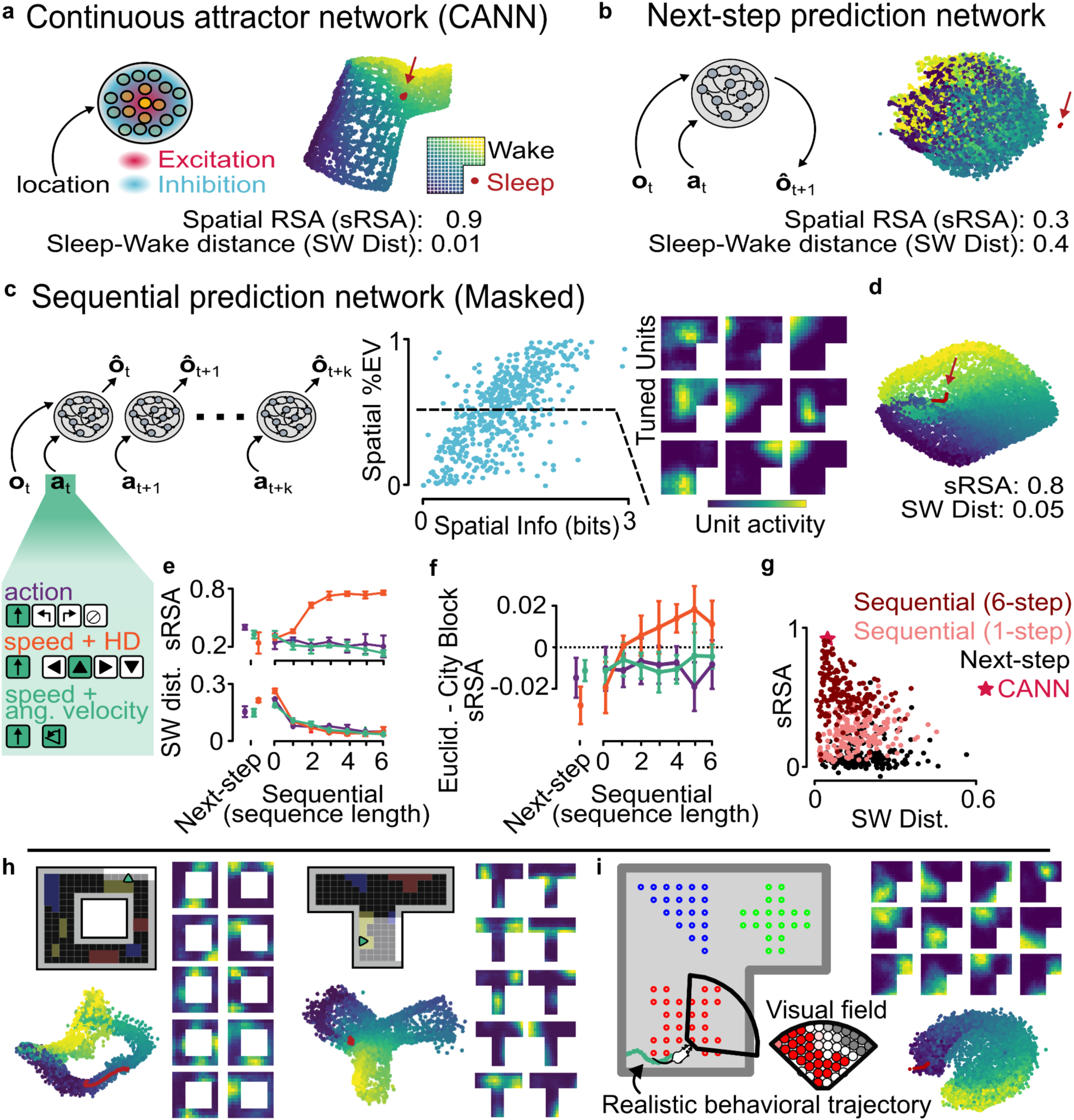
A continuous attractor-based representation emerges from sequential predictive learning. **a:** Isomap visualization of population activity in a continuous attractor network (CANN), in which units assigned to represent nearby positions are connected with recurrent excitation and those representing distant positions are mutually inhibiting. Each point represents a population activity vector at a point in time, colored by the position of the agent (wake), with sleep timepoints in red. Spatial representational similarity analysis (sRSA) measures the correlation between spatial and neural activity distances, while Sleep-Wake distance (SW Dist) measures the neural distance between sleep and wake activity in the network. **b:** Similar to A, but for the Next-Step pRNN from Figure 1. **c:** In the masked sequential predictive network, predictions (*ô*) are made for a k-length sequence of held-out sensory observations at the current timestep (Note that after timestep t+k, the network will receive observation and action corresponding to t+k+1; see also **Methods** and **Figure S1**). (Right) Spatial tuning properties of recurrent units from an example masked predictive learning with a sequence length of k=5, trained with actions encoded as speed (0 or 1, for forward motion) and head direction. **d:** Similar to A, for the k=5 masked network from panel C. **e:** sRSA and SW Dist for next-step pRNNs and masked pRNNs with k=0 (autoencoder), k=1 (single-step), and k=2-6 (sequential), each trained with one hot (action), speed+HD, and speed+AV encoding of the action input, **at**. Mean and standard deviation calculated over 9 seeds. **f:** Difference between sRSA calculated using Euclidean vs city-block distance, for all networks in **e**. Higher Euclidean sRSA indicates a euclidean, rather than behavioral-step-based map of space. **g:** sRSA and SW Dist for a hyperparameter-matched population of Next-step, single-step, and 5-step masked pRNNs. **h:** Neural manifolds and example spatial tuning curves for a 5-step masked pRNN trained in environments with different geometries. **i:** Neural manifolds and example spatial tuning curves for a 5-step masked pRNN trained in a continuous environment ^80^ (speed continuous).

We thus wondered whether other forms of predictive learning might produce an attractor-based representation of the environment. Recent findings indicate that recurrent autoencoders can also contain spatially-tuned cells ^77^, and masked predictive learning, an approach in which an autoencoder is trained to predict inputs at held-out timesteps, has been found to produce good internal models in e.g. vision and language tasks ^78^. Under a masked prediction paradigm, visual inputs can be masked for multiple time-steps (**Figure 3c, S1**), which we hypothesized would encourage the recurrent connectivity to maintain a coherent representation of the agent’s current state and capture transitions through space. Indeed, we found that multi-step, or sequential, predictive learning led to the emergence of a neural manifold that reflected the layout of the environment (sRSA=0.8+/-0.05) and constrained neural activity during sleep (SW Dist=0.05+/-0.02, i.e. is an attractor, **Figure 3d**, **S9**). The presence of this manifold supported the use of this representation to plan an effective route to a remembered location in the environment (**Figure S10**), or navigate to a goal in the face of obstacles (**Figure S10**). We will thus refer to this kind of representation as a “cognitive map” ^1,79^.

Interestingly, despite the ability of all masked networks to solve the prediction task and develop spatially tuned cells, only sequential networks trained with speed and head direction (HD) formed a cognitive map (**Figure 3e, f**). Specifically, an attractive manifold (low SW Dist) only emerged in networks trained to predict at least two time-steps of masked observations (**Figure 3e**) and the mapping to space (high sRSA) didn’t emerge in one-step masked networks or in sequential networks receiving action identity or speed and angular velocity (**Figure 3e, Figure S11**). Furthermore, sequential predictive networks trained with speed and HD information were the only networks in which their spatial representation more closely reflected Euclidean distance than the number of steps between locations (**Figure 3f**). These results were robust across networks with different hyperparameters (**Figure 3g, S12**), consistent across environments with different geometries (**Figure 3h**), and in a continuous environment with realistic behavioral trajectories^80^ including those with different movement statistics (**Figure 3i, Figure S13**).

While the use of layer normalization, dropout, and noise injection (see Methods) improved the stability of training and offline activity, and spatial tuning in the next-step RNNs (**Figure S14**), it did not promote the formation of a continuous attractor in the next step networks, and their presence wasn’t necessary for its emergence in sequential RNNs (**Figure S14**).

How can the next-step predictive RNN have spatially tuned representations but not an attractive neural manifold that maps the spatial layout of the environment? This apparent contradiction was a result of the network’s representation of space being superseded by strong representations of action identity and visual inputs (**Figure S15**), which were only weakly represented in the sequential predictive network (**Figure S15**). Interestingly, increasing the degree of visual ambiguity in the environment reduced the spatial representation in next step, single step, and action identity encoded networks, but not in sequential net with speed+HD action encoding (**Figure S16**), further supporting the idea that sequential predictive learning with speed and head direction information encourages a cognitive map solution. The appearance of a cognitive map manifold could be attributed to the prevalence of cells that have more of their variance explained by spatial position. Despite having tuning curves with the same amount of spatial information, units in the sequential prediction networks had more of the variance in their activity explained by spatial position (**Figure S17**), and more of these spatially-tuned cells had unimodal tuning curves for single spatial locations (**Figure S17**). Across the hyperparameter population, sRSA was correlated with the fraction of tuned cells rather than mean SI (**Figure S17**).

However, the prevalence of spatially tuned units in a network does not necessarily imply that all cells show place cell-like tuning (i.e. single place field rate maps). Indeed, a careful examination of unit response properties (**Methods**) revealed a diversity of spatial tuning properties (**Figure 4a**), including border cells, single-field cells with a diversity of field widths (**Figure 4d**), and a majority of cells showing complex or conjunctive tuning (**Figure 4c**). We hypothesized that the connectivity structure of the hippocampus (specifically, the sparse recurrent connectivity with lognormally-distributed excitatory weights in CA3^66^) is a key factor in determining its spatial tuning. To test this, we initialized the network with sparse connectivity and lognormal weights, and used an optimization procedure that preserves skewed weight distributions^81^ (**Figure 4b, Methods).** We found that this increased the prevalence of single-field cells in the network (the proportion of single-field cells increased from 19±2% to 30±1% among all cells in the network, and from 38±3% to 65±2% among spatially-tuned cells, **Figure 4c**) and increased the field widths (**Figure 4d**). This connectivity profile did not only affect the tuning of single units, but also improved population coding. Specifically, sparse-logNormal connectivity decorrelated the population vectors between distant locations (**Figure 4e**), in-line with recent experimental results ^21^.

**Figure 4:**
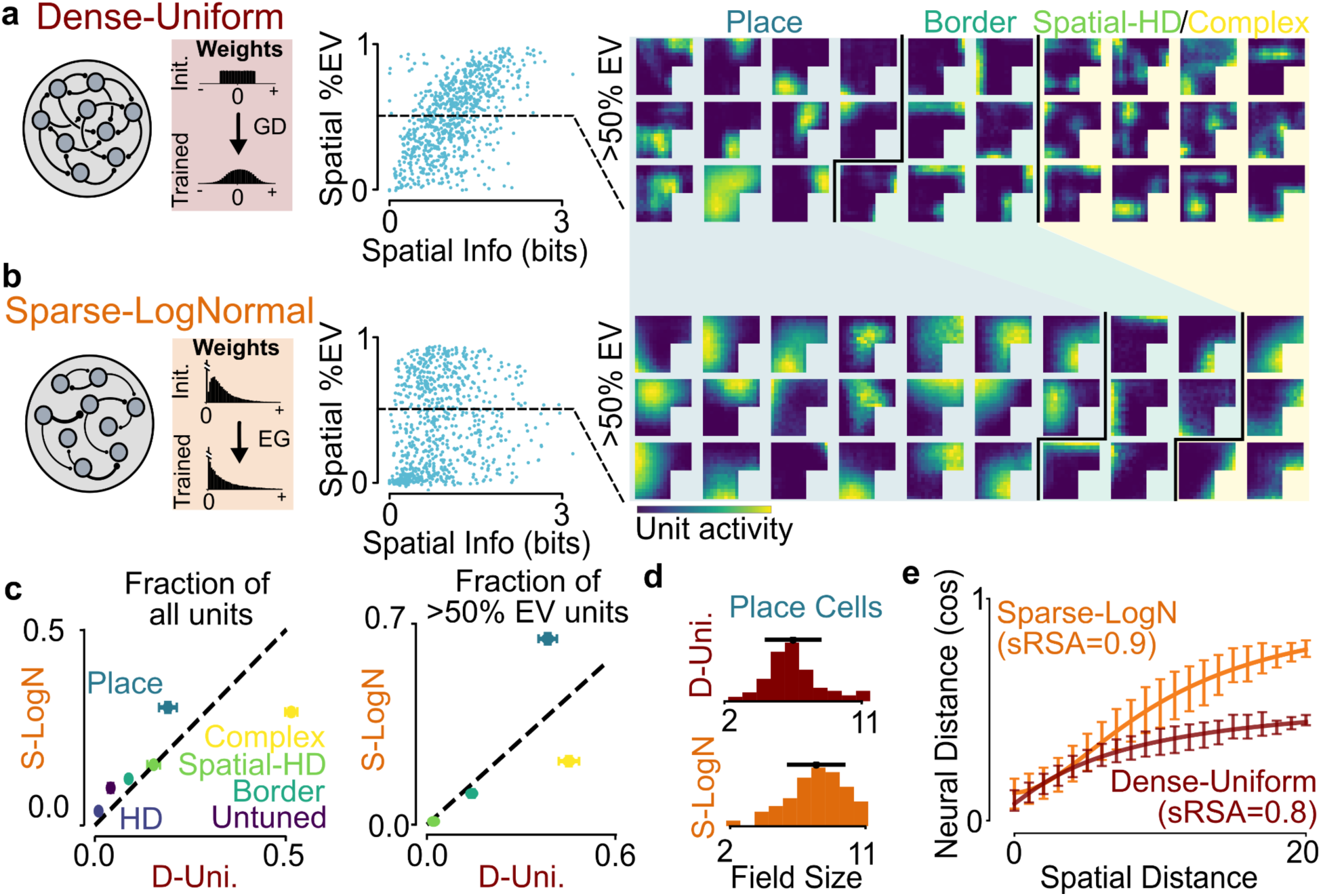
Sparse lognormal connectivity leads to an orthogonalized manifold with more single-field cells. **a:** Standard “vanilla” RNNs (dense-uniform), like those in the rest of the paper, are fully-connected networks, and trained with gradient descent (GD) which does not preserve Dale’s law. (Middle) Spatial information and explained variance by position for all units in a sequential masked (k=5) network. (Right) A random selection of spatial tuning curves in the network from cells with more than 50% of their activity variance explained by position, separated into cells classified as place cells, border cells, and conjunctive or complex cells (see Methods for classification). **b:** Sequential masked (k=5) networks were initialized with sparse (5%) recurrent connectivity with lognormally distributed weights, and trained with an exponentiated variant of gradient descent (EG) that preserves skewed distributions ^81^(see Methods). (Middle/Right) Compare to panel a, for the sparse-Lognormal network. **c:** Proportion of each cell type in the dense-uniform and sparse-logNormal networks (mean+/-std over 6 seeds). Note the sparse-logNormal network has proportionally many more place cells, at the expense of complex cells. (Left) All cells. (Right) Restricted to cells with >50% of their variance explained by spatial position. **d:** Distribution of field sizes for place cells in the sparse-logNormal and dense-uniform networks. Bar indicates mean +/- standard deviation. **e:** Mean neural distance between population vectors at all pairs of timepoints, as a function of the spatial distance between the agent’s location at those timepoints. Error bars indicate mean +/- std, fit to Hill equation with range of equilibrium neural distances shown over 6 networks with different seeds (see Methods).

While the pRNN is trained to predict upcoming sensory inputs, the hippocampus does not receive direct sensory signals. Instead, hippocampal inputs consist of high-level representation in the entorhinal cortex, for example in the form of grid cells. To ensure that our results did not depend on the choice of input, we trained networks with grid cell-like inputs (**Figure S18**). Like networks that learned from visual input, these networks formed a cognitive map representation of the environment (**Figure S18**), with single-field cells whose prevalence was increased by sparse-lognormal recurrent connectivity (**Figure S18**).

We next asked if the network could learn to represent multiple environments (**Figure 5a**). After training on up to 7 environments, networks were able to predict observations in each environment without a context cue, and networks had comparable values of sRSA as networks trained on a single environment (compare to **Figure 4e**). One possibility is that the same cognitive map was transferred across environments. Another possibility is that, like in the hippocampus ^82,83^, the network generated separated maps that supported prediction in each environment. Isomap projection of population states revealed environment-specific and non-overlapping population states (**Figure 5b**), which were linearly separable (**Figure 5c**). Learning more maps increased the dimensionality of network activity, from a 6-dimensional (intrinsic/nonlinear^84^) manifold embedded in a 13-dimensional (embedding/linear) space for a single environment to a representation with an intrinsic dimensionality of 11 embedded in a 36-dimensional space for a network that learned 7 environments (**Figure 5d-f**). This demonstrates that sequential predictive learning creates a high dimensional representation that can independently map multiple environments to different manifolds, similar to the hippocampus.

**Figure 5:**
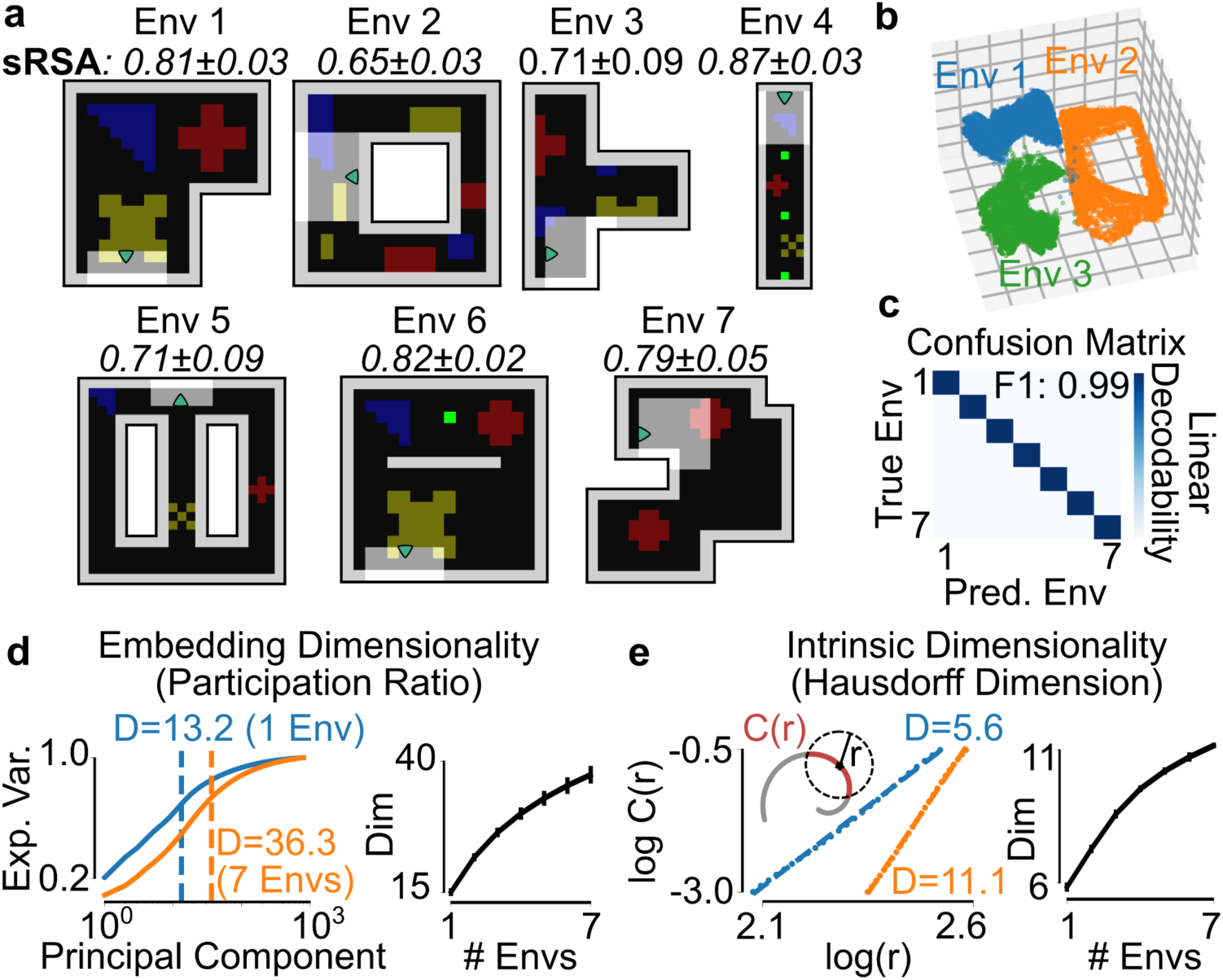
Sequential predictive learning creates a high dimensional representation that can independently map multiple environments. **a:** 7 gridworld environments in which sequential predictive RNNs (masked k-5, with sparse-logNormal connectivity) were trained. sRSA for each environment calculated from networks trained on all 7 environments, mean+/-std over 5 random seeds. **b: I**somap visualization of population activity of a network trained in all seven environments, in environments 1-3. **c:** Confusion matrix of a linear decoder trained to decode environment identity from population activity. **d**: Embedding (linear) dimensionality of network activity. (Left) Cumulative explained variance of the principal components of network activity for networks trained on 1 and 7 environments. Dimensionality is calculated as the participation ratio of the eigenvectors. (Right) Embedding dimensionality of networks trained on 1-7 environments. Error bars reflect mean+/-std over 5 seeds. **e**: Intrinsic (nonlinear) dimensionality of network activity. (Left) Correlation integral C(r) (the fraction of pairs of points within radius r) of network activity for networks trained on 1 and 7 environments. Dimensionality is calculated as the slope of log C(r) versus log r. (Right) Intrinsic dimensionality of networks trained on 1-7 environments. Error bars reflect mean+/-std over 5 seeds.

In summary, sequential predictive learning of egocentric sensory observations, using an agent’s speed and head direction, leads to the emergence of a population code for space with CANN-like attractor properties.

### Offline activity in sequential predictive RNNs can autonomously replay recent locations or follow a head direction query

In contrast to the next-step predictive networks (**Figure 2**), offline activity in the sequential predictive RNNs maintained a coherent representation of position in the absence of sensory input. When the network was driven only by internal noise, the decoded position at each timestep was spatially concentrated (**Figure 6a, b**), and made local transitions around a single location which was different for each sleep epoch (**Figure 6c**, **Figure S19**).

**Figure 6:**
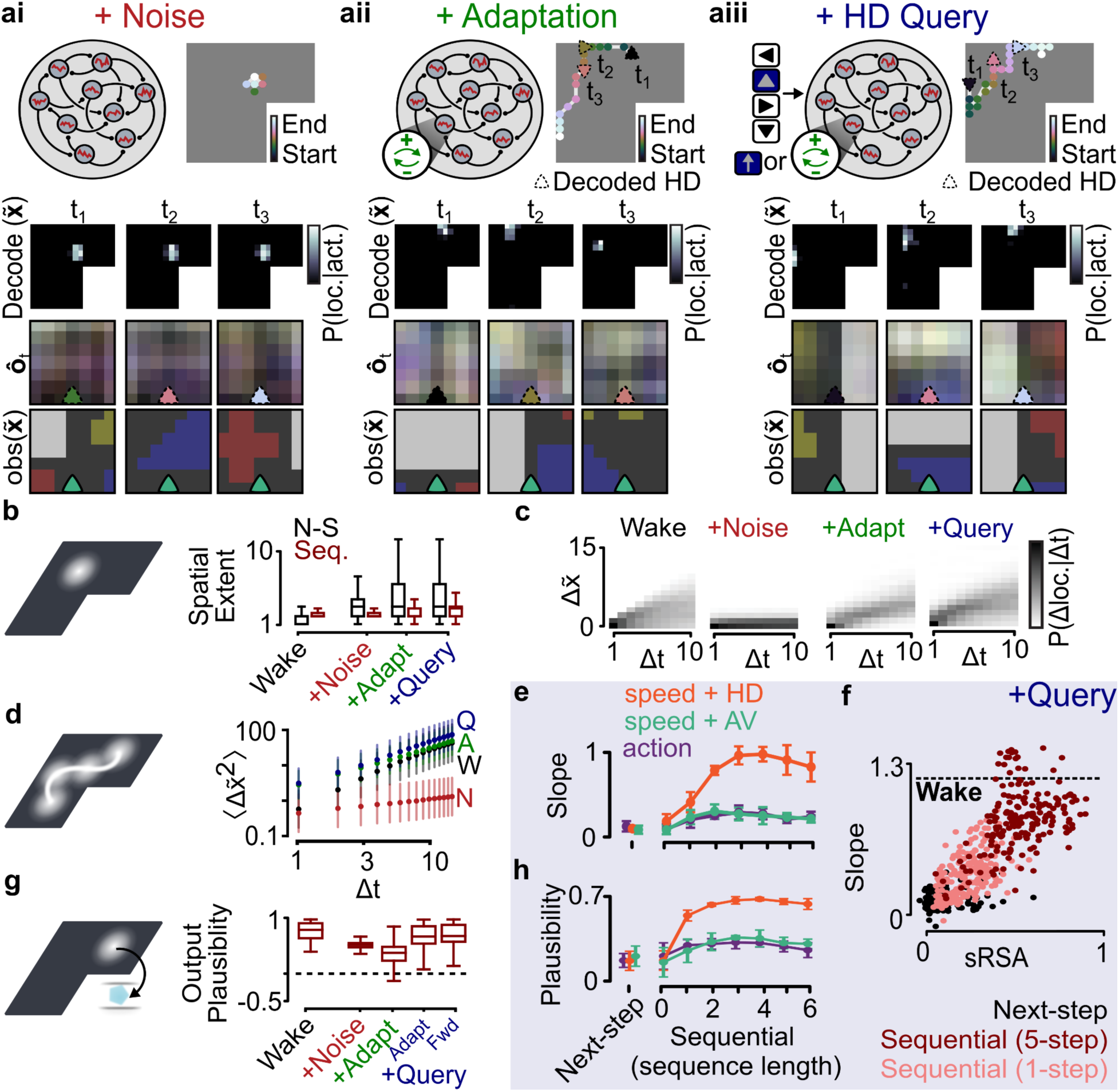
Sequential predictive RNNs generate offline trajectories that can follow a head direction query. **a:** Example subset of a sleep trajectory from the example masked pRNN with a sequence length of k=5, including decoded posterior, observation output, and true observation from the decoded viewpoint. Similar to Figure 2f. Offline activity (sleep) is generated in the absence of sensory input using internal noise (**i**), the addition of a neuron-autonomous negative feedback (adaptation, **ii**), and a virtual head direction query with either adaptation or virtual speed query (**iii**). **b:** Distribution of spatial extent of the decoded posterior for offline activity in the next-step and sequential (k=5) masked pRNNs, with wake and each sleep mechanism. Compare to Figure 2g. **c,d:** Spatial diffusion statistics of the decoded trajectory during wake and each sleep mechanism. Compare to Figure 2i. **e:** log(<**Δx**^2^>)/log(**Δ**t) slope for query-generated offline activity in next-step pRNNs and masked pRNNs with k=0 (autoencoder), k=1 (single-step), and k=2-6 (sequential), each trained with one hot (action), speed and head direction (speed+HD), or speed and angular velocity (speed+AV) encoding of the action input. Mean and standard deviation calculated over 9 seeds. **f:** log(<**Δx**^2^>)/log(**Δ**t) slope as a function of sRSA for a hyperparameter-matched population of next-step, single-step, and 5-step masked pRNNs. **g:** Distribution of output plausibility of the decoded posterior for offline activity in the sequential (k=5) masked pRNNs, with wake and each sleep mechanism. Compare to Figure 2h. **h:** Output plausibility for query-generated offline activity in next-step pRNNs and masked pRNNs with k=0 (autoencoder), k=1 (single-step), and k=2-6 (sequential), each trained with one hot (action), speed+HD, or speed+AV encoding of the action input.

Offline activity in the hippocampus is not restricted to single locations, though, shows extended paths that travel through the environment and often recapitulate salient or rewarded positions from recent exploration ^25^. A common way to produce trajectories in CANNs and other hippocampal models is the addition of adaptation ^46,85–87^ (**Figure 6a**), i.e. slow negative feedback on each neurons’ activity which is thought to play a key role in producing the dynamics of offline activity in the hippocampus ^88^. We found that adding an adaptive variable to our RNN units resulted in extended sleep trajectories (**Figure 6c, S19, S20**), with similar statistics to wake trajectories (**Figure 6d**). Remarkably, in addition to forward-moving transitions, decoded offline trajectories included diagonal and 2-step transitions, “reverse” transitions in the opposite direction of the decoded head direction (**Figure S19**), and previously-unseen trajectories across the two branches of a T-shaped maze that were not in the training data (**Figure S19**, compare to ^28^). Thus, the network can autonomously produce generative replay that reflects the structure of the environment, but is not strictly limited to the observed, behaviorally-constrained state transitions during wake. Increasing the learning rate during a waking trial caused the network to shift from generating plausible but random trajectories, to recapitulating locations visited within the trial during the subsequent sleep epoch (**Figure S21**).

In addition to replaying recent locations, it’s been suggested that offline hippocampal activity could be used for directed planning ^10,30^ and imagination ^13^. Motivated by observations that the neural circuits that represent head direction are spontaneously active during offline activity in the brain ^89^, we hypothesized that the same input that indicated the agent’s actions during wake could direct trajectories during sleep (**Figure 6a**). Indeed, like adaptation, the addition of an action “query” produced wake-like transition statistics (**Figure 6c,d, S20**), by influencing the direction of replay transitions (**Figure S19**). This behavior was only seen in the sequential predictive networks trained with speed and head direction information (**Figure 6e**), and was related to their ability to form a cognitive map as measured by sRSA and S-W distance (**Figure 6f**). Interestingly, the ability to produce wake-like trajectories only relied on the head direction component of the action signal, while either fictitious speed or adaptation could modulate the velocity of this movement (**Figure S19**). In addition, the head direction query improved the plausibility of visual generation, producing output that mimicked the associated sensory inputs at the replayed locations during sleep (**Figure 6g,h, S20**). Together, these results reveal that sequential predictive learning produces networks containing a cognitive map that achieves the three desiderata for offline activity (**Figure 1c**), namely it: (1) maintains a coherent representation in the absence of input, (2) produces extended trajectories that can replay recent experiences or directed via a head direction query, and (3) can recapitulate the learned sensory associations at the represented positions.

### Rollout-based sequential predictive learning produces theta sweep representational dynamics and rapid learning

While the learning procedure above recapitulates several experimental observations, it requires a large amount of sensory “data” – more precisely, a map is only learned after about 2000 trials of 500 steps each (**Figure S23**). In contrast, a neural manifold emerges in the hippocampus after only a few days of exposure to a novel environment ^21,23^ and the hippocampus is able to rapidly form associations between positions and novel sensory information ^90^. Further, unlike the hippocampus ^58,59^, the masked approach described above (**Figure 3**) predicts held-out present, rather than future, sensory observations. Once the prediction sequence is completed (after k timesteps), the network receives the observation corresponding to the k+1 step in the environment. In self-supervised ^91^ and reinforcement learning ^92^, sequential prediction is often learned via “rollouts”: prediction of multiple future timesteps, which results in higher data efficiency ^93^. Therefore, we modified the learning procedure to use a rollout-based approach in which the RNN predicts multi-step sequences of future observations (t+1:t+k) every timestep before it receives the next time-step’s sensory inputs (t+1, **Figure 7a, S1**).

**Figure 7:**
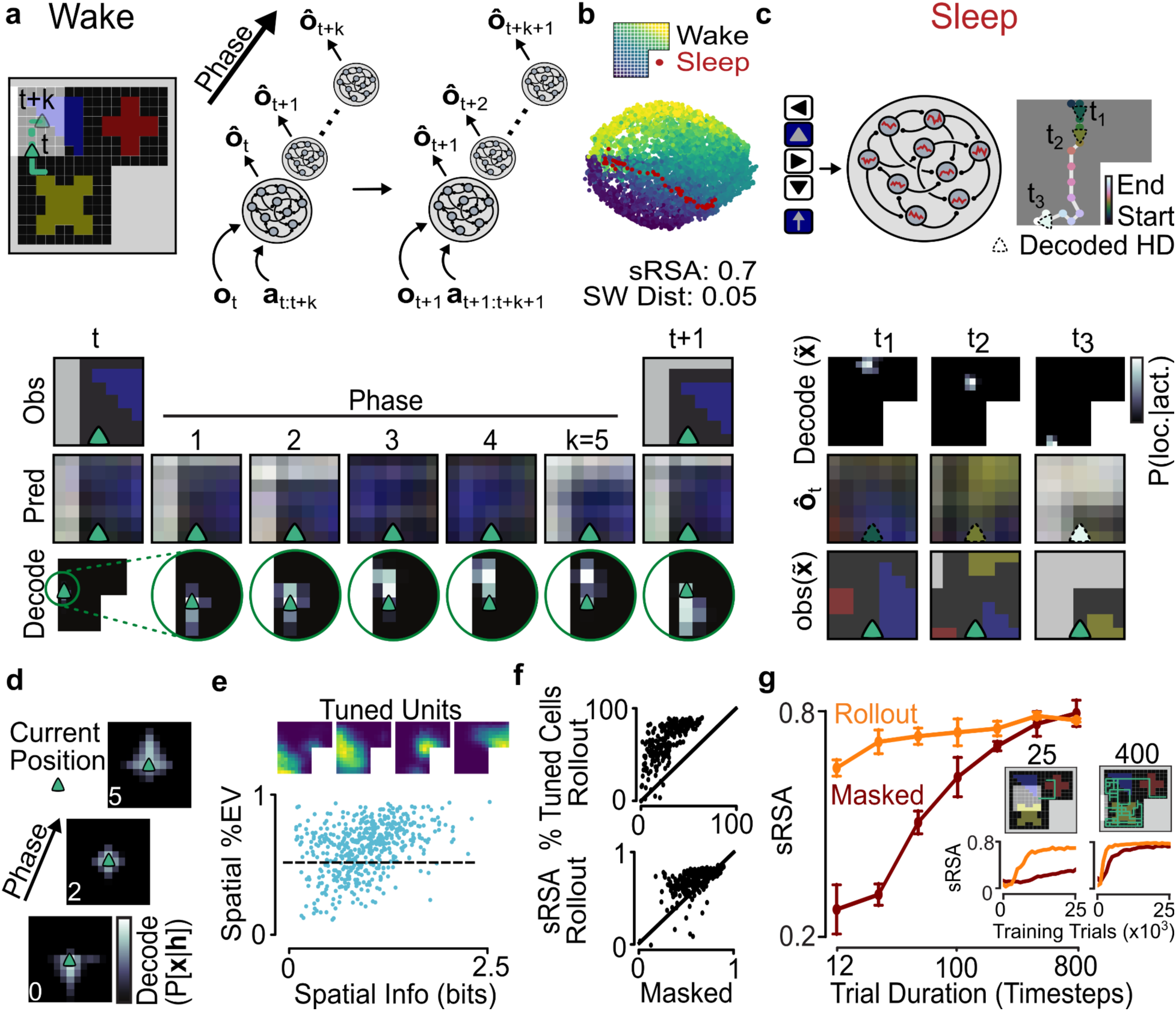
Rollout-based sequential predictive learning produces theta sweep representational dynamics and rapid learning. **a:** In the rollout-based predictive network, predictions (*ô*) are made for a k-length sequence of future sensory observations, based on the current sensory observation and the next *k* actions. Unlike the masked approach in Figure 3, the rollout-based network only steps forward in the world after predicting k timesteps; after the rollout it receives the observation corresponding to timestep t+1, and the next k actions (See also **Figure S1**). **b:** Isomap visualization of the neural activity manifold for an example rollout-based network with a sequence duration of *k* = 5. Similar to Figure 3a. **c:** Example query-generated offline activity in the rollout-based sequential pRNN. Similar to Figure 6Aiii. **d:** Mean decoded posterior relative to the agent’s current position for activity at early, middle, and late phases of the prediction cycle. **e:** Example units, and spatial tuning properties of all network cells in the rollout-based sequential pRNN. **f:** Percentage of tuned cells (top) and sRSA (bottom) for a population of hyperparameter-matched rollout-based and masked pRNNs with *k* = 5. **g:** sRSA for rollout-based and masked sequential networks (*k* = 5), trained using trials of different durations. Training for all networks was run for 5×10^4^ trials. Mean and standard deviation calculated over 9 seeds. (Inset) sRSA over the first 2.5×10^4^ trials for networks trained with 25 and 400-step trials.

Like sequential predictive networks using the masked approach, this rollout-based predictive RNN learned a continuous attractor map (**Figure 7b, S22**), which maintained a coherent spatial representation during sleep and produced wake-like trajectories with plausible sensory associations (**Figure 7c**). Interestingly, when the decoder was applied to successive phases of the rollout, the position represented by the network would “sweep” ahead of the agent as it moved through the environment (**Figure 7a,d**). This dynamic representation was strikingly similar to hippocampal theta sweeps: spiking sequences representing locations that project ahead of the animal every ∼120 ms, coordinated by an 8Hz theta oscillation^94^. To learn the attractor map, the network had to be provided with the prospective sequence of future speed and head direction (**Figure S21)**, akin to having planned its actions in advance. However, once the map was learned, the network was able to simulate possible but untaken future trajectories during rollouts, when provided with a hypothetical action sequence (**Figure S22**). Intriguingly, the sweeping representations began immediately behind the agent, despite the RNN being trained to predict present and future, but not past, observations. As also noted in ^95^, this was because the decoder was trained to detect the agent’s current location using neural activity from the entire rollout cycle, analogous to the experimental practice of decoding in ∼120ms bins (one theta cycle), and thus was trained on timesteps in which the network was largely predicting the agent’s immediate future observations. The resulting decoder was thus biased to decode positions slightly before the predicted observation, resulting in a backward shift of decoded position when applied to individual timesteps of the rollout.

Rollout learning drastically improved learning performance and the emergence of a cognitive map. Rollout networks had more spatially-tuned cells (**Figure 7e-f**), developed a higher-sRSA representation of space than their masked counterparts (**Figure 7f, Figure S23**), and were better at sensory prediction (**Figure S23**). Further, this approach was able to more-rapidly form new sensory associations when a novel object was introduced in the environment, which persisted in the network’s output even when the object was removed **(Figure S24**). The reason for this improvement is that the network was more data efficient - it was able to form a continuous attractor map with fewer trials of shorter sequence duration (**Figure 7g, Figure S23**).

Altogether, we found that rollout-based sequential predictive learning improved the speed and robustness of learning continuous attractor maps, and replicated the experimentally observed patterns of theta sweep “lookaheads”. Hence, our findings suggest that theta sequences are a signature of sequential predictive learning in the hippocampus.

### Sequential predictive learning produces task-relevant non-spatial representations

In addition to space, the hippocampus has been reported to represent non-spatial variables when they’re relevant to a task or behavioral context. For example, in a spatial alternation task, cells have tuning curves that are specific to the future behavior, i.e. whether the animal will turn left or right ^16,17^. Likewise, in a multi lap task where animals receive a reward after 4 laps around a circuit, cells are selective for different laps ^18^. In a tone-cued behavior task, cells develop selectivity for space, tone frequency, and distance to goal, which together form a manifold that captures relevant information for action plans ^19^. Our model was able to recapitulate each of these results when trained with a task guided action policy (**Figure 8**).

**Figure 8:**
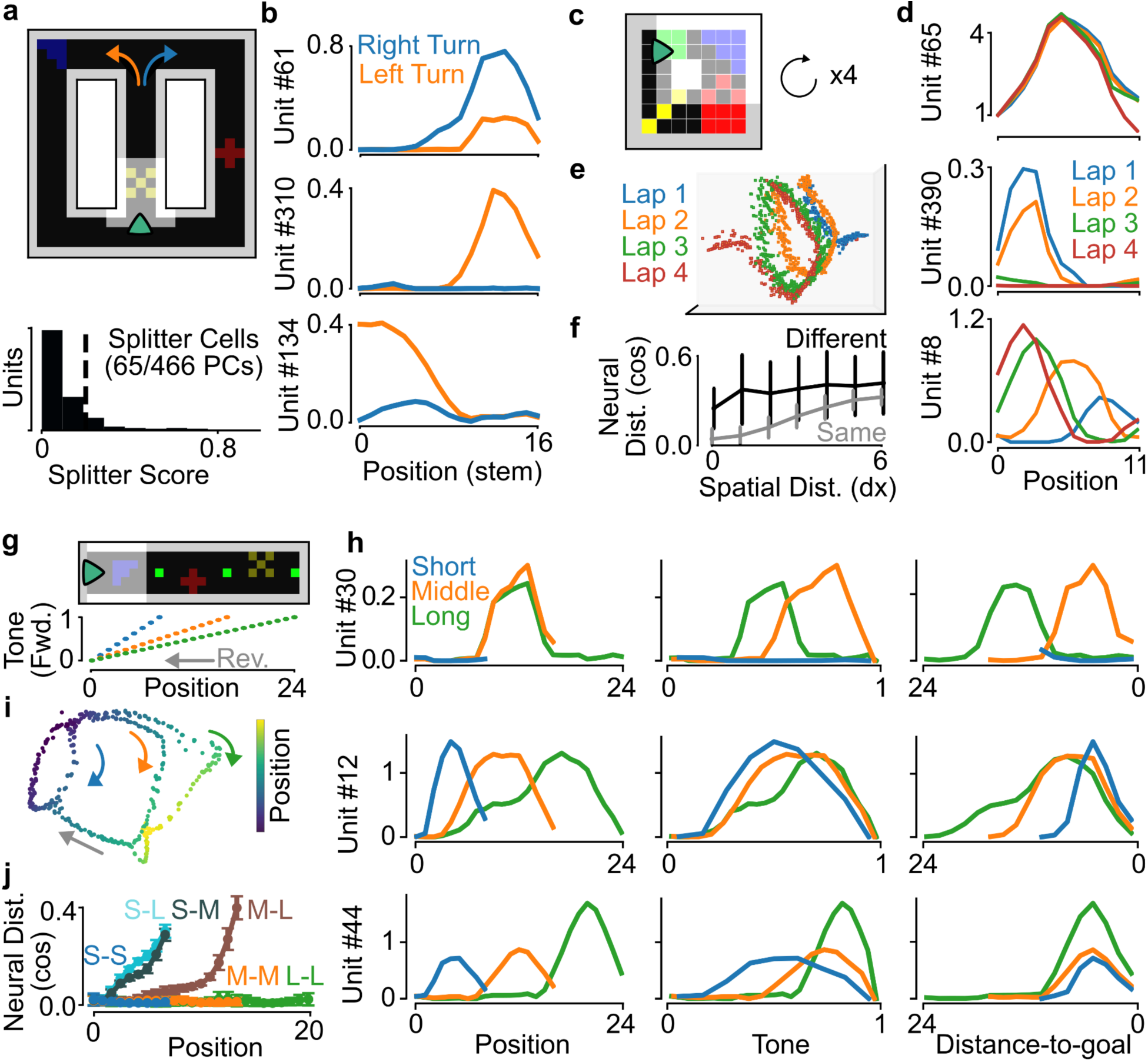
Sequential predictive learning produces task-relevant non-spatial representations. **a:** The rollout-based network was trained in a figure-8 maze (compare to ^16,17^), using a policy that caused the agent to alternate between left (orange) and right (blue) turns. (Bottom) Distribution of splitter scores (see Methods) of units in the trained network. **b.** Spatial tuning curves along the middle stem, separated into trials when the agent was going to turn left or right, calculated from the activity of three example units in the network. **c:** The rollout-based network was trained in a circular maze (compare to ^18^), using a policy that caused the agent to traverse the circle for four consecutive laps before pausing and receiving a “reward” stimulus (see Methods). **d:** Spatial tuning curves of the linearized position around the maze, separated into each lap, calculated from the activity of three example units in the network. **e:** Isomap projection of the population activity on the maze, colored by lab identity. **f:** Mean +/- standard deviation of neural distance between population vectors as a function of the linearized spatial distance between pairs of time points, and divided into timepoints when the agent was on the same lap or different lap around the maze. **g:** The rollout-based network was trained on a linear maze with goal positions at three different distances along the maze (compare to ^19^). As the agent traversed the maze in the forward direction, a “tone” stimulus was input to the network, which increased at one of three different gains (indicating short, middle, and long trials). The agent followed a policy by which it turned around at the goal position corresponding to the maximum tone. **h:** Tuning curves calculated from the activity of three different units, with respect to the agent’s position, the value of the tone, and the distance to the goal, and divided into short, middle, and long trials. **i:** Isomap projection of the population activity on the maze, colored by the agent’s position. Note the distinct arms for short, middle, and long trials, indicating a different neural representation on each trial trial type for the same position. **j:** Mean +/- standard deviation of neural distance between population vectors between different trial types at each position on the maze. Note that same trial types (S-S, M-M, L-L) have a very similar neural population representation at the same position, while different trial types (S-L, S-M, M-L) have a distinct neural representation (increased neural distance) at the same position.

First, we trained the sequential predictive model using an action policy that caused the agent to alternate between left and right turns in a figure-8 maze (**Figure 8a**, top), similar to the experiments of ^16,17^. This led to the emergence of many “splitter cells” (**Figure 8a**, bottom), whose spatial selectivity was determined by the upcoming turn that the agent would make, e.g. firing more before a right turn than a left turn or vice versa (**Figure 8b**).

Next, we trained the sequential predictive model using an action policy that had the agent go around a doughnut maze four times before a novel “reward stimulus” was presented (**Figure 8c**), after which the agent would wait for 10 time-steps before proceeding to the next trial, similar to the experiments of ^18^. Training in this environment produced cells in the recurrent network that were standard place cells (**Figure 8d**, top), but also notably, place cells that were selective for the lap (**Figure 8d**, middle) and cells whose place field moved as a function of lap number (**Figure 8d**, bottom). This resulted in a population manifold that tracked lap number (**Figure 8e**), such that distance in neural space reflected both distance in space and lap number (**Figure 8f**). In addition, we found that some neurons in this environment developed temporal selectivity during the waiting period between each trial (**Figure S25**), similar to the “time cells” reported in the literature ^20^, and that place field locations tended to shifted against the direction of movement when the agent switched from unbiased exploration to directionally-biased behavior (clockwise lap action policy, **Figure S25**), similar to the backwards shift observed along often-traveled trajectories ^96^.

Finally, we trained the network in a linear track environment in which a simulated audio tone would increase in pitch until the network arrived at one of three goal locations, which were either a short, middle, or long distance from the starting location (**Figure 8g**), similar to the experiments of ^19^. In this situation the sequential predictive model developed cells that were selective for spatial position (**Figure 8h**, top), tone frequency (independent of location; **Figure 8h**, middle), and distance to the upcoming goal (independent of location or tone; **Figure 8h**, bottom). Again, this resulted in a task-relevant population manifold that had distinct “arms” reflecting the different trial types and progression through the trial (**Figure 8i**). As a result, the angle between representations for the same position on different trial types reflected the difference between the trial types (short, medium, long; **Figure 8j**). Altogether these results demonstrate that if an agent is engaged in task-relevant behaviour then sequential predictive learning will represent task-relevant variables, including non-spatial variables, as seen repeatedly in the hippocampus.

## Discussion

In this study, we have used a RNN model to demonstrate that sequential predictive learning provides a unifying theory for hippocampal representation and replay. In line with previous work ^55,56,77^, we found that learning to predict sensory inputs robustly leads to the development of spatially tuned cells in recurrent neural networks, and that this can produce a representation of the environment on a low-dimensional neural manifold ^55,56^. However, we found that the presence of spatially tuned cells is not *sufficient* to guarantee a cognitive map with the ability to generate offline replay. Nonetheless, a continuous attractor manifold consistently emerged in networks trained to predict multi-step sequences of sensory inputs using speed and head direction information, which allowed the networks to simulate plausible trajectories offline. This offline “replay” activity emerged from spontaneous activity in the network, without the need for an explicit replay algorithm, and could recapitulate trial positions, generate novel trajectories, or be directed by a head direction query. We found that the efficiency of cognitive map formation was increased by interleaving predictions of future sensory inputs with each action in the environment, which produced representations that mimicked theta sweeps observed in the hippocampus. These results show that multi-step sequence prediction is a promising unified account of hippocampal operations.

Our work builds upon previous studies that have explored the relationship between predictive models and spatial representations in the hippocampal-entorhinal system. This past work has shown that: (1) Aspects of spatial tuning in the hippocampus are well-explained by predictive representations ^39,57^, which can be learned from spatially tuned inputs with predictive Hebbian learning ^97^, TD learning ^98^, or spike-timing dependent plasticity ^98–100^; (2) Learning to predict memory embeddings from actions can link allocentric and egocentric representations ^101^; (3) Prediction can link spatially tuned inputs and relational structures ^102^; (4) Cloned hidden-Markov models that use discrete states to predict observations from actions can recapitulate many of the features of the hippocampus and support offline evaluations ^103,104^; (5) Engaging in path integration (i.e. predicting spatial locations from sequences of actions) leads to the emergence of grid cells and continuous attractor dynamics ^49,51^; (6) Training recurrent networks to predict hippocampal spiking data or linear place cell sequences can reproduce various features of hippocampal activity, including spike cross-correlations and sequence replay ^105^. Beyond these predictive models, past work has shown that spatial representation characteristic of the medial entorhinal cortex (grid cells and toroidal manifolds) emerge in networks trained on path integration or spatial classification losses ^49,50,71,72^, though, it should be noted that these approaches require spatial position information either in their inputs or their training objective and do not produce single-field place cells or replay. Our work demonstrates that the findings from the models listed above can be brought together under the umbrella of sequential predictive learning of sensory data in RNNs, applied to sensory observations that are rich, spatially ambiguous, and redundant. Critically, this same objective also gives rise to hippocampal place cells, continuous attractor dynamics, directed and autonomous replay of novel trajectories (c.f. ^28^), and non-spatial population manifolds that reflect task structure (**Figure 8**), properties that are not simultaneously captured by any of these alternative approaches.

More broadly, sequential predictive learning unifies the ideas underlying three distinct views of hippocampal operations. In addition to connecting views that the hippocampus is a predictive map ^39^ with those that the hippocampus implements a CANN ^41^, sequential predictive learning aligns with views that the hippocampus is a sequence generator ^40,62,64,104^. These views are based on extensive physiological observations of theta sequences ^94^, which are tied to behavioral steps such as whisking ^106^, footsteps ^107^, sniffing ^108^, or saccades ^109^, and are essential for subsequent replay ^110–112^. They are further supported by recent models demonstrating that associating observations with sequences of discrete states can reproduce many physiological phenomena associated with the hippocampus ^104^, and classic models demonstrating that sequence learning in neural networks can reproduce many of its psychological functions ^62^.

One of the most surprising results in our study was that spatially tuned cells are not a guarantee of a low-dimensional neural manifold with topological correspondence to the environment, or cognitive map. Supporting this distinction between spatial tuning and a cognitive map, experimental work has shown that, while place cells appear almost instantaneously in novel environments ^113^, a cognitive map only emerges after multiple days of exposure ^21,23^ and relies on weakly tuned cells ^23,52^. Further, our observation that predicting sequences of sensory inputs was sufficient for map formation and replay, but not spatial tuning, mirrors physiological results that place cells emerge in the hippocampus before sequential activity during development ^114,115^ and that perturbing its sequential structure disrupts its ability to produce replay, but not its associative code for space ^112^.

Once formed, this learned cognitive map supports a spectrum of offline activity without any modification to the recurrent connectivity. Driven by noise alone, the attractor maintains a spatially coherent representation of individual locations in the absence of sensory input (**Figure 4a-i, 4b**), resembling the reactivation of single positions during hippocampal sharp-wave ripples ^112^. The addition of neural adaptation (a well-established mechanism for generating sequential dynamics in attractor networks ^85,86^) converts this stationary reactivation into autonomously-generated trajectories across the learned manifold (**Figure 4c**, **S19**). Because the network has learned a continuous spatial representation rather than a discrete set of experienced state transitions, these trajectories can include novel paths, such as diagonal and reverse movements, that were never present in the training data, consistent with experimental observations of replay of never-taken shortcut routes ^28^, and is biased towards the positions of recent, high-learning-rate trials. A virtual head direction input further enabled the network to direct replay along specific trajectories (**Figure 4g**), and output associated sensory information at the replayed positions. Thus, the contribution of sequential predictive learning is not any individual replay mechanism, but rather a flexible, generative map that can support reactivation, autonomous replay, and directed replay through the same learned representation.

In addition to predicting sequences of sensory inputs, our results suggest that a head direction signal may play a key role in learning a cognitive map from viewpoint-dependent observations. While the inclusion of a head direction signal is, admittedly, allocentric, it has a striking resemblance to the inputs that convey an animal’s head-direction and forward velocity to the hippocampal system ^116^, direct theta sweeps ^117^, and emerge prior to place cells during development ^118^. Indeed, where previous studies used one-hot encoded actions or cartesian velocities to learn spatial representations, our results show that these less-physiological encodings do not facilitate learning a representation that supports a cognitive map, despite their ability to learn the prediction task and develop spatially-tuned cells. Further, the head-direction system conveys a coherent signal during sleep which is disconnected from the animal’s actual head direction ^89^ and shows increased activity immediately prior to hippocampal replay ^119^. While adaptation alone was sufficient to produce replay trajectories in our model, a virtual head direction input enabled the network to output associated sensory information at the replayed positions, and was able to influence the direction of replay trajectories. Together, these results predict that head direction is not necessary for forming place cells, but that is instrumental in the formation of a cognitive map, the production of replay that recapitulates learned associations for downstream processing ^37^, and can support the use of replay for directed planning and imagination. This critical role of the head direction system is an experimental prediction that could be tested in future work.

There are a number of limitations in this work that should be considered. First, we trained our networks from a random initial state, which is not an accurate reflection of learning in the hippocampus. There is substantial evidence showing that the hippocampus has pre-existing connectivity structures on which learning operates ^66,120^, and a rich anatomically-constrained circuit structure ^121–123^. One prominent hypothesis is that these structures endow the hippocampus with a reservoir “library” of existing sequences ^66,67,124^, which can be associated with arbitrary sequences of sensory input. We hypothesize that such an initialization would facilitate rapid, and potentially even single-shot, sequential predictive learning. Moreover, any adult animal will have ample experience with many environments, and this previous knowledge would be brought to bear on learning “novel” environments ^125^. Future work could explore how sequential prediction behaves in networks with structured initializations and/or training on multiple continuously learned environments.

Second, while we were inspired by theta dynamics for the sequential learning algorithm, we did not concern ourselves here with the biological realism of the synaptic plasticity mechanism. We instead aimed to show that theta sweeps may be a signature of sequential predictive learning, and could be used as data-efficient rollouts predicting the sequence of future sensory input. However, the rollout-based predictive network can potentially be implemented by biologically-plausible plasticity mechanisms. One possibility is that behavioral time-scale plasticity (BTSP) driven by plateau potentials ^126^ could provide the eligibility traces required to maintain a record of past predictions and compare them with current observations in dendritic compartments with spatiotemporally-segregated inputs ^127^. While BTSP has generally been studied in feedforward inputs from CA3 to CA1 ^128^, they can also potentiate the recurrent synapses necessary for sequential prediction in CA3 ^129^. Another possibility is spike timing-dependent plasticity (STDP). Theta-timescale sequences place successively active place cells within the temporal window for synaptic potentiation via STDP ^130,131^. Disruption of theta-timescale sequential structure impairs the development of predictive spatial representations ^112^, and the oscillatory dynamics of the hippocampus restrict inputs to different phases of the theta oscillation ^132^, which could effectively act as the masking mechanism for sensory inputs ^100^.

The ability to both represent space and generate replay from high-dimensional egocentric sensory inputs is, we believe, a significant advance of our model. Previous models of replay have tended to rely on reduced representations that presuppose spatial tuning (either in the input ^47,87,111^, local connectivity ^46,47,87,133^, or learning target ^51^), or treat replay as a simple recapitulation of specific spike sequences in a one-dimensional environment ^87,134,135^. However, it should be noted that unlike our model, the hippocampus does not receive direct visual inputs, but instead receives signals via the entorhinal cortex that are processed by multiple cortical structures, and that place cells can form in the absence of visual inputs (i.e. in complete darkness ^136^). While we did demonstrate that our results hold when the predicted input is a grid cell-like representation of space, the medial entorhinal cortex provides highly organized inputs to the hippocampus, including additional specialized neuronal populations such as landmark cells that make it possible to rapidly encode new environments and may be critical for learning more complex environments like the multi-sensory real world. We anticipate that the emergence of spatial tuning does not rely on the specific use of visual inputs in the prediction, but will emerge so long as a network learns to use actions to predict any sensory modality during movement, and that a cognitive map will emerge when those sensory inputs are sequences. Moreover, a cognitive map should also form when the predictions are on higher-order representations of sensory inputs, rather than raw sensory information.

Finally, we lack a formal account of how multi-step predictive learning leads to the emergence of manifold-based cognitive maps and the capacity for replay, which we leave for future work. While previous work has explained the emergence of attractors from connectivity ^49,137,138^, the question of how a specific representation emerges from a given loss is a critical unsolved problem in theoretical neuroscience and beyond the scope of this work. Conceptually, we suspect that multi-step prediction changes the nature of the learning problem. In essence, one-step prediction is often just an action-dependent transformation of the current inputs (see e.g. **Figure S14**), though it does require some place conditioning to deal with ambiguous and redundant locations, hence the emergence of place cells. In contrast, multi-step prediction leads to a combinatorial explosion of the possible transformations of the inputs, and the formation of a map becomes the most optimal way to achieve accurate predictions. Another way of saying this is that predicting what will be seen multiple steps into the future requires the network to have an internal representation of the different locations in the environment and *how they relate to each other*. If it does not, then the network has no way of predicting accurately multiple steps into the future given the explosion of possible inputs. As a result, using recurrent connections for sequential predictive learning forces the network to learn the latent generative structure of the observations ^65^. Finally, we hypothesize that the inclusion in the training data of timesteps without observation inputs (e.g. masked timesteps or rollouts), renders observation-free (i.e. sleep) activity “in-distribution” for the network.

Our model allows us to make several explicit predictions that can be tested with existing experimental techniques. First, disrupting head direction inputs to the hippocampus (for example, by silencing the para- or post-subicular projection ^139^) should impair the formation of a spatial manifold and the generation of directed replay, while leaving single-cell spatial tuning largely intact. Second, disrupting theta-timescale sequential structure during exploration should impair the formation of a manifold representation, in addition to the previously-reported impairment of replay generation ^112^. More specifically, our model predicts that theta sequences are not merely a readout of the cognitive map, but are instrumental in its formation. Third, if head direction signals serve as “queries” that direct hippocampal replay, then head direction activity in the para- or postsubiculum should precede and predict the trajectory of a larger subset of replay events than expected by chance. Simultaneous recordings from the postsubiculum and hippocampus could test whether replay direction aligns with the para- or post-subicular head direction representation, and whether silencing this input during sleep modifies the statistics of replay.

In conclusion, our study demonstrates the ability of sequential predictive learning to account for hippocampal representation during active exploration and replay during behavioral quiescence, from egocentric information available to an animal. This represents, we believe, a major step forward in our understanding of the algorithmic-level understanding of the hippocampus, and provides support to the theory that a major algorithmic operating principle for the hippocampus is that of sequence prediction.

## Methods

### Data Availability

No experimental data was used in this manuscript.

### Environment

#### Gridworld

The gridworld environment was developed using the minigrid package (https://minigrid.farama.org/, Chevalier-Boisvert et al 2023). An 18×18 L-shaped room (**Figure 1B**) was generated using walls and colored floor tiles, which each corresponded to a unique [R, G, B] color value. Walls ([0.6, 0.6, 0.6]) were impassible. Floor ([0.3, 0.3, 0.3]) and colored floor tiles ([0.45, 0.45, 0], [0, 0, 0.45], and [0.45, 0, 0]) were passable, and arranged in different shapes on the floor to serve as visual cues.

For each “wake” trial, the agent was initialized at time *t* = 0 at a random position, ***x***_0_ and head direction, ***HD***_0_, in the environment. It then took a series of *T* actions, which were randomly selected from (move forward, rotate left, rotate right, stop) with probability 0.6, 0.15, 0.15, and 0.1, respectively. As the agent moved, the observation at time *t* corresponded to the 7×7×3 image of tiles immediately in front of it, with the agent’s current position at the bottom center of the image with color [0.5, 0.3, 0.3] (**Figure 1A**). This image was then flattened, resulting in *T* + 1 observation vectors, ***o****_t_* ∈ ℝ*^Nobs^* = *obs*(***x****_t_*, ***HD****_t_*), *N_obs_* = 147, which started with the initial observation at *t* = 0 and ended with the observation following the final action, ***a****_T_*.

The actions, ***a****_t_*, were encoded in one of three different forms: 1) a *one-hot encoding* vector of length *N_act_* = 4, in which the action at time t was represented by a 1 and the other actions were represented by a 0; 2) a *speed and angular velocity encoding* vector of length *N_act_* = 2, in which the first element was 1 if the agent took the move forward action and 0 otherwise, and the second element was 1 if the agent took the rotate right action and -1 if the agent took the rotate left action; 3) a *speed and head direction encoding* vector of length *N_act_* = 5, in which the first element was 1 if the agent took the move forward action and 0 otherwise, and the other 4 elements were a one-hot encoding of the agent’s current head direction (North, South, East, West).

#### Rat in a Box

The continuous environment was developed with the RatInABox toolkit (https://github.com/RatInABox-Lab/RatInABox, George et al 2024). For comparability, the room was designed as close as possible to the one described above (**Figure 3i, S6a**): a 1×1m room of the same L-shaped form with three distinct classes of objects positioned to fit the locations of colored floor tiles in the Gridworld environment.

Agent’s movements in the RatInABox are generated using two independent Ornstein-Uhlenbeck processes for rotational velocity and linear speed as described in the original paper. The agent was initialized with the following parameters: time discretization (the amount of time a single step takes, *dt*): 0.1s; speed_coherence_time (time over which linear speed decoheres): 0.7s; speed_mean: 0.2m/s; rotational_velocity_coherence_time (time over which rotational speed decoheres): 0.08s; rotational_velocity_std: 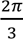 thigmotaxis (tendency for agents to linger near walls [0 = not at all, 1 = max]): 0.2; wall_repel_distance: 0.1m.

The observations along the movement trajectories were collected using the Field of View (FoV) neurons feature. Each of these neurons has preferred distance from the agent and angle relative to agent’s heading. If the object of interest is within proximity of this preferred location, the neuron outputs a value according to a double-gaussian model, with a maximum of 1. The agent was initialized with separate sets of FoV neurons for each of the three classes of objects and for the walls: {*F*_1_, *F*_2_, *F*_3_, *F*_w_}. Each set contained 48 cells that tiled a sector in front of the agent with a radius of 0.33m and the angle boundaries (-45°,45°) relative to the agent’s head direction. Thus, each neuron for one class of object had corresponding neurons with the same preferred distance and angle for each other object class and for the walls. Note that having orthogonal representations for each type of object and for the walls is different from what was done in the Gridworld environment where the floor marks and the walls shared the RGB channels (for example, both red and yellow mark were present in the red channel, and the walls were represented in all three channels). To match the Gridworld RGB pattern, we transformed the outputs from FoV neurons into a 48×3 matrix where 48 is the number of neurons and 3 is for red, green, and blue channels. *F_1_* output was added to channel 1 (red), *F_2_* output was added to channel 3 (blue), *F_3_* output was added to channels 1 and 2 (green), and *F_w_* output was multiplied by *0.39* and added to all three channels. This matrix was then flattened, resulting in observation vector *o_t_* of size *N_obs_* = 144. The actions *a_t_* were encoded by a vector of length *N_act_* = 13 with first element corresponding to the agent’s linear speed, and the rest one-hot encoding its head direction that was binned into 12 sectors.

Alternatively, we used grid cells activity as the observational input (**Figure S18**) to the model using the GridCells feature of the package. Grids are modeled as the rectified sum of three cosine waves at 60 degrees to each other. We specified the scales (wavelengths) and orientations for 10 modules, each containing 15 cells with different phase offsets drawn randomly from a uniform distribution. Below is the list of pairs of scales and corresponding orientations:

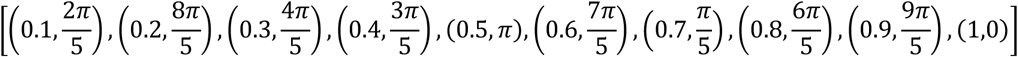

Resulting observation vector *o_t_* was of size *N_obs_* = 150. The actions were encoded same as for visual observations.

### Predictive RNNs

The core of each predictive RNNs followed the same general architecture.

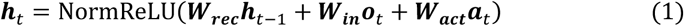

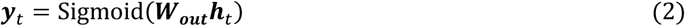

With activation functions,

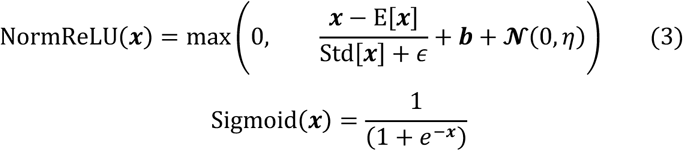

In which ***h****_t_* ∈ ℝ*^N^* is the state of the network at time t, ***b*** ∈ ℝ*^N^* is a neuron-specific bias, ***W_rec_*** is the (*N*x*N*) recurrent weight matrix, 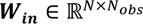 and 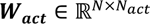 are the input weight matrices for observation and action, respectively, 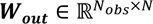 is output weight matrix, *ε* = 10^−4^ is a small constant to prevent division by 0, and *η* is the standard deviation of uncorrelated noise injection to the network (Lim et al 2021, Krishna et al 2024). The first term in the second argument of equation 3 corresponds to layer normalization (Ba et al 2016), which was replaced with ***x*** for networks trained without layer normalization (**Figure S14**). For networks trained without noise injection (**Figure S14**), *η* was set to 0.

The three different pRNN architectures (Next Step, Masked, Rollout) were distinguished by the timesteps in which ***o****_t_* was available to the network and the identity of the observation predicted by the output, ***y****_t_*, per the loss function, ℒ.

In the *next-step* architecture, the network received observation input at each timestep, and was trained to match the output to the subsequent observation,

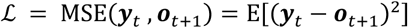

i.e., ***y****_t_* = ô*_t_*_+1_, where ***o***i indicates a prediction of variable ***o*.**

In the *masked* architecture, the network was trained to match the output to the current observation,

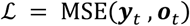

i.e., ***y****_t_* = ô*_t_*. However, the observation was only provided as input to the network every *k* + 1 timesteps. That is,

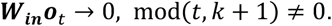

Like the masked architecture, the *rollout* architecture was trained to match the output to the current observation. However, at every observation timestep, *t*, the RNN would rollout *k* timesteps,

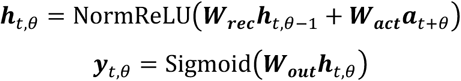

Where ***h****_t_*_,0_ = ***h****_t_*, *θ* ∈ 1: *k* indexes the phase of the rollout, and the network was trained to match the output to the corresponding present (*θ* = 0) or future (*θ* > 0) observation.

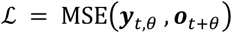

i.e., ***y****_t_*_,*θ*_ = ô*_t_*_+_*_θ_*. For rollout-based networks trained with “looped” time, ***h****_t_*_–1_ = ***h****_t_*_–1,0_ in eqn. 1, for “continuous” time, ***h****_t_*_–1_ = ***h****_t_*_–1,*k*_.

#### Training

The trainable network parameters Ω = (***W_rec_***, ***W_in_***, ***W_act_***, ***W_out_***, ***b***) were initialized as follows:

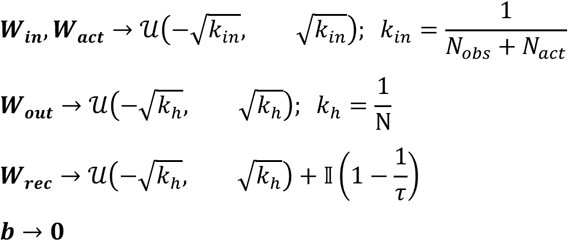

Where *τ* is a hyperparameter that sets the neural timescale. ***o****_t_* was passed through a dropout layer. The networks were trained using backpropagation through time (BPTT) with the RMSProp algorithm as implemented in *pytorch* (Paszke et al 2019), with *α* = 0.95 and *ε* = 10^−7^, for 8×10^4^ single-batch trials in the environment, unless otherwise stated. Network state was randomly initialized from **N**(0, *η*) on each trial, and the data for each trial was collected from the environment, as specified in “Environment – Gridworld”, above. The learning rate was set by a global learning rate hyperparameter, *λ*, and then further scaled for each trainable parameter by its initialization value or an additional hyperparameter, as follows:

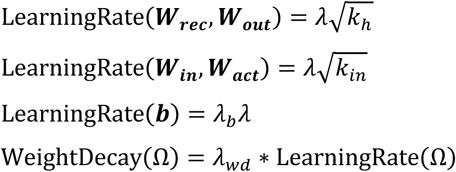

The training hyperparameters can be found in Table 1 for the example networks used and the range for the random hyperparameter sweep.

**Table 1.**
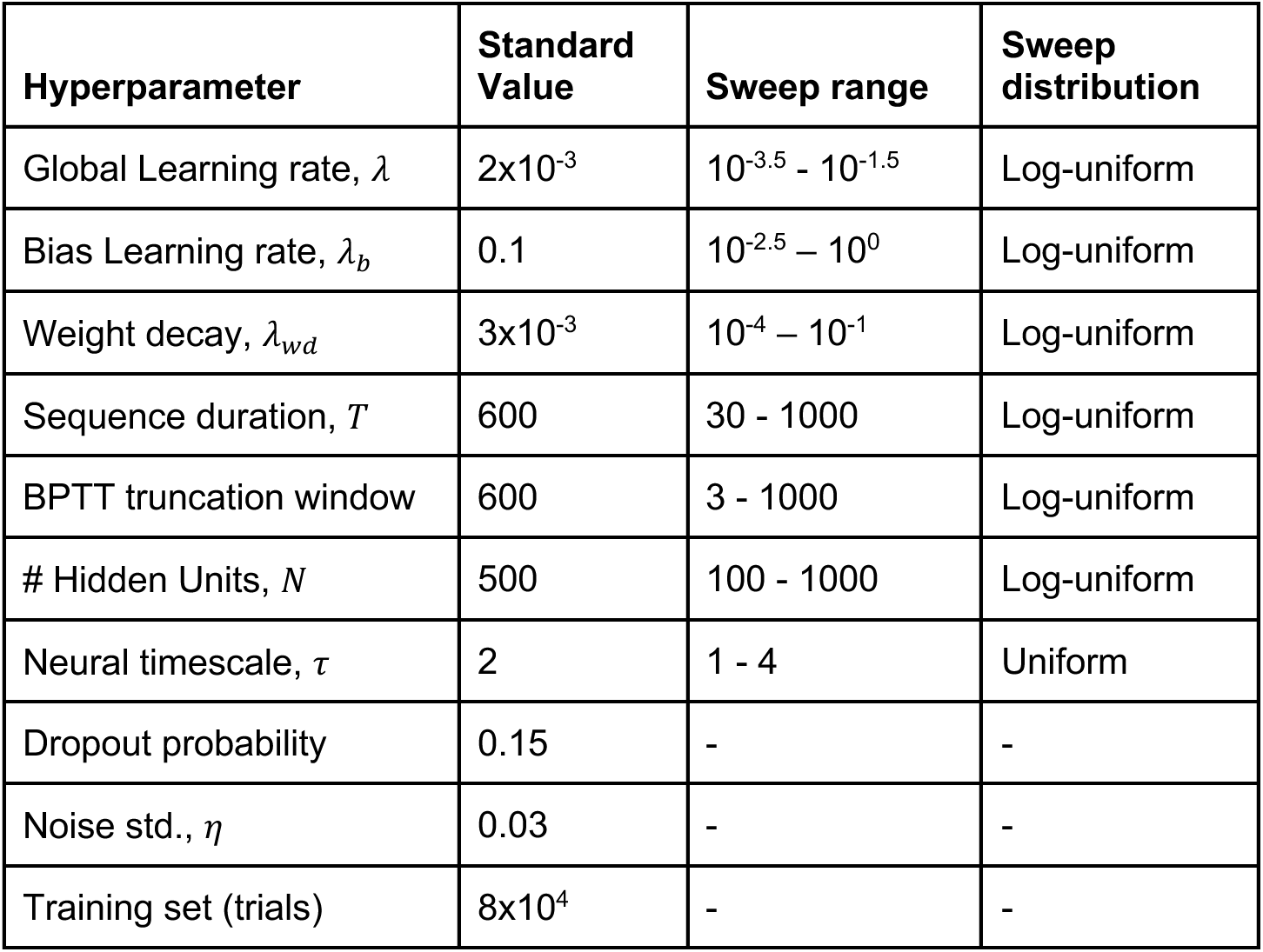
Training hyperparameters. Standard value was used in all experiments, unless stated otherwise. Values for the random hyperparameter population (**Figures S7, S12**) were sampled from the sweep range using the sweep distribution.

To train networks on multiple environments (**Figure 5**), trials from all environments were randomly selected and interleaved in a minibatch of 32 trials. All other training parameters were kept the same. Networks were trained for 140 epochs total, 1000 gradient steps/epoch) across five random seeds for every number of environments used.

#### Sparse-lognormal network

The sparse-lognormal networks were initialized with ***N*** = **800** hidden units, as follows:

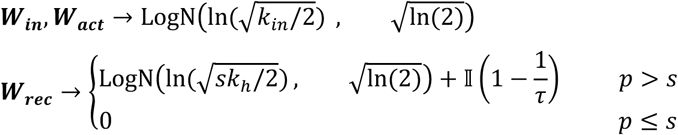

Where LogN(*μ*, *σ*) is a random variable drawn from a lognormal distribution where *μ* and *σ* are parameters specifying the mean and standard deviation of the underlying normal distribution (as specified in the pytorch builtin function `tensor.log_normal_`, producing a weight distribution with standard deviation matching that used for the dense-uniform networks), *p* is a random variable drawn from U(0, 1), and *s* = 0.05 is the sparsity of the network. The network was trained using the RMS prop algorithm with exponentiated gradient descent (Cornford et al 2025) for the lognormal parameters (***W_in_***, ***W_act_***, ***W_rec_***) with learning rate *λ*_E*G*_ = 2×10^−3^ and weight decay *λ*_w*d*,E*G*_ = 1×10^−8^ and compared to “vanilla” RNNs (dense-uniform, similar to all other networks described) with the same number of units (**Figure 4**).

### Sleep activity generation

To generate offline activity (sleep) in each RNN, the network was initialized from ***N***(**0**, ***η***), and allowed to run for ***T_sleep_*** timesteps (as specified for each analysis) without the observation input,

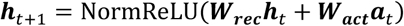

This resulted in a timeseries of neuronal activations, ***h***_0:*T*_*_sleep_*, that were “spontaneously generated”, that is, produced without reference to input from the environment and relying only on the recurrent weight matrix ***W_rec_*,** a random network initialization, gaussian noise, and potentially a fictitious action sequence, ***a***_1:*T*_*_sleep_* (see below). Critically, like in the brain, the position represented by the network is only available via a decoder trained to decode the agent’s position from the network’s distributed activity as it explores the environment during wake, which in this case corresponds to the probabilistic linear decoder designated below.

In the case of noise-generated offline activity, the action was also removed (***a****_t_* = 0), and the standard deviation of noise injection (Eqn. 3), *η*, was increased to 0.2 unless otherwise indicated (**Figures S8, S19**).

In the case of adaptation-generated offline activity, the action was also removed (***a****_t_* = 0), an additional cell-autonomous slow variable, ***c****_t_* ∈ ℝ*^N^*, was added to the RNN equations (compare to eqns 1, 3),

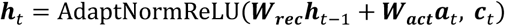

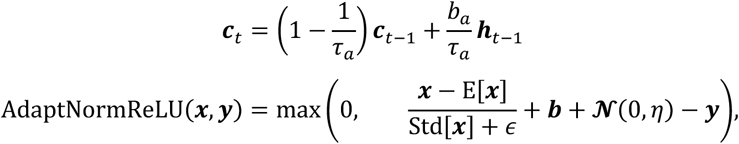

where *τ_a_* = 100 and *b_a_* = 0.5 are the timescale and strength of adaptation. The effect of modifying their values is shown in **Figure S20**.

In the case of query-generated offline activity, ***a****_t_* was randomly generated following the same action probabilities as wakefulness. In the case of the speed and head direction encoding, the dependence on the speed signal was tested by setting the speed component to 0 and using the network with adaptation (**Figure S20**).

### Analysis

#### Spatial representation

To decode the position represented by the network, the agent was run for a trial (see Environment – Gridworld) of duration *T* = 1.5 × 10^4^ timesteps, and the sequence of hidden unit activity, ***h****_t_*, was used to train a linear decoder with the following architecture.

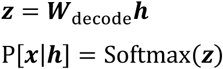

in which P[***x***|***h***] is the posterior probability of the agent being in position ***x*** given a hidden unit activity vector, and 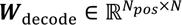 is the decoding matrix with *N_pos_* being the number of possible positions in the environment and *N* being the number of hidden units in the network.

***W***_decode_ was trained to minimize the following loss:

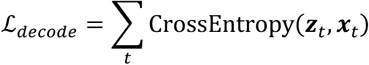

in which 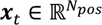, is a one-hot encoding of the agent’s position at time *t*. The network was trained using the AdamW optimizer with learning rate = 10^-3^ and weight decay = 3×10^-1^, for 5×10^3^ weight updates, with a batch of 5×10^3^ random timepoints used in each update to prevent overfitting. To decode the head direction to calculate the expected viewpoint during sleep, a second decoder was trained with ***x****_t_* = ***HD****_t_* as a one-hot encoding of the agent’s head direction at time *t*.

The decoding error was calculated by running an additional test trial of duration *T* = 2 × 10^3^ timesteps, and calculating the cityblock distance between the agents actual and decoded position,

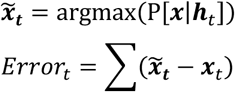

To calculate spatial tuning curves and spatial information, the agent was run for a trial (see Environment – Gridworld) of duration *T* = 1.5 × 10^4^ timesteps. The tuning curve, or mean activation as a function of position, ℎ*_i_*(*x*) = *E*[ℎ*_i_*|*x*] was calculated for each unit using the *pynapple* package for neural data analysis (Viejo et al., 2023). The spatial information was then calculated for each unit’s tuning curve (Skaggs et al., 1992),

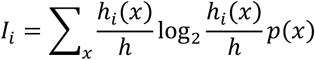

Where ℎ is the unit’s mean activation. This metric is a standard metric of spatial tuning curves, and measures its entropy.

To calculate the spatial explained variance (%EV_S_) for each unit, the agent was run for another trial (see Environment – Gridworld) of duration *T* = 5 × 10^3^ timesteps. %EV_S_ was taken to be the decrease in the variance of the unit’s activity when the expected rate given the spatial tuning curve was removed. That is,

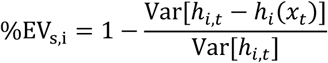

#### Cell type classification

**To further classify units based on their spatial tuning properties (Figure 4), we calculated the following additional metrics for each cell, using the cell’s tuning curve for space or head direction:**

##### Number of place fields

To determine the number of place fields, the 2D autocorrelation of each unit’s tuning curve was calculated using scipy’s `correlate2d` function. The number of fields was then taken to be mod(num_peaks+1,2), where num_peaks is the number of peaks in the autocorrelation greater than 15% of the max and of size bigger than 3 squares.

##### Place field size

The place field size was calculated as the square root of the number of squares in a contiguous area with the autocorrelation’s central peak, whose value was greater than 50% of the peak.

##### Place field asymmetry

The place field asymmetry was calculated by fitting an ellipse around the squares in a contiguous area with the autocorrelation’s central peak, whose value was greater than 50% of the peak. The asymmetry was taken to be the ratio of the major and minor axis of that ellipse.

##### Border score

The border score was calculated as 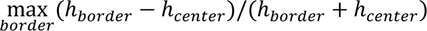 where (ℎ*_border_* is the mean rate of the unit along each border of the environment, and ℎ*_center_* is the mean rate of the unit in all squares that are not within 2 squares of any border.

##### Head direction information

Head direction information was calculated using the same equation as spatial information, but with a unit’s tuning curve for head direction instead of space.

Cells were then grouped based on the following procedure: first, cells whose EVs was less than 15%, and with spatial information and head direction information was less than 0.5 bits were classified as “untuned”. Cells whose EVs was less than 15%, and with spatial information less than, but head direction information greater than 0.5 bits, were classified as “HD cells”. Of the remaining cells, those with border score greater than 0 and asymmetry greater than 3 were classified as “border cells”. Of the remaning cells, those with a single place field and EVs greater than 50% were classified as “place cells”. Of the remaining cells, those with spatial and HD information greater than 0.5 bits were classified as “spatial-HD” and the remaining cells were classified as “complex”.

#### Replay analysis

To analyze the spatial extent, output plausibility, and trajectory statistics of the represented activity during sleep, the networks was run for 15 sleep trials of *T_sleep_* = 300 timesteps each, and the linear decoder was used to calculate the decoded posterior, P[***x***|***h****_t_*], and decoded position *x̃_t_* and head direction 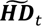 at each timestep.

Spatial extent of each timepoint was taken to be the furthest distance at which the 2D autocorrelation dropped below 0.01.

Plausibility of each timepoint was taken to be the spearman correlation between the network output ***y****_t_* and the environmental observation from the decoded viewpoint, 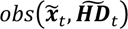.

Trajectory statistics were calculated as in (Stella et al., 2019). The average squared Euclidian distance, 〈Δ*x̃*^2^〉, was calculated between the decoded positions of all pairs of timepoints separated by Δ*t* = [0: 15] timesteps in the same trial, where [*x*: *y*] indicates the list of integer values between x and y. A linear fit was then calculated for the relationship between log〈Δ*x̃*^2^〉 and log〈Δ*t*〉, producing a slope, *α*, that quantified the degree to which the trajectories reflect sub or supra-diffusive motion, and an intercept, *G*, that reflected the average speed of trajectory propagation.

#### Neural manifold analysis

To calculate the manifold properties of each network, we simulated a *T* = 1.5 × 10^4^ timestep wake trial and a *T_sleep_* = 1000 timestep sleep trial (see Environment – Gridworld).

To visualize the neural manifolds during wake and sleep, we used Isomap (Tenenbaum et al., 2000) with the hidden unit activation from 4×10^3^ randomly sampled timepoints from the wake trial and all timepoints from the sleep trial. The *sci-kit learn* implementation of isomap was used with parameters n_neighbors=150, n_components=2, and metric=’cosine’.

Spatial RSA (sRSA) was taken to be *ρ*m*D*_E_(***x****_t_*, ***x****_t_*_′_), *D_C_*(***h****_t_*, ***h****_t_*_′_)n, where *ρ*(*X*, *Y*) is the spearman rank correlation between X and Y, 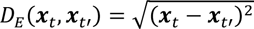 is the Euclidian distance between the agent’s position at all pairs of timepoints *t* and *t*′ during the wake trial, and 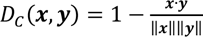 is the cosine distance between the hidden unit activations at all corresponding pairs of timepoints ***h****_t_* and ***h****_t_*_′_. To quantify the relationship between neural distance in the network and spatial distance in the environment (**Figure 4**), the relationship between *d**x*** = *D*_E_(***x****_t_*, ***x****_t_*_′_) and *d**h*** = *D_C_*(***h****_t_*, ***h****_t_*_′_) was fit to a Hill equation,

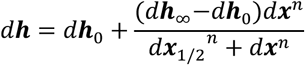

Where *d**h***_0_, *d**h***_∞_, *d**x***_1/2_, and *n* parameterize the average neural distance at the same position, the average neural distance at maximal spatial distance, and the midpoint and steepness of the transition between these two extremes.

Sleep-Wake distance (SW dist) was taken to be the average distance between the neural activations during the sleep trial and the closest point in neural space during the wake trial. That is,

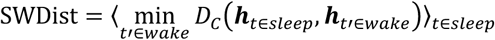

To analyze the attractiveness of the representational manifold (Figure S9), we first generated pRNN hidden states that were uncharacteristic of its typical wake states. For that, we performed PCA (n_components=100) over 6×10^5^ steps of wake activity and projected this activity to the resulting low-dimensional space, yielding coefficients *z*t. We then computed the empirical radius for each embedded point

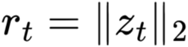

and set a typical radius *r_typ_* at the 90^th^ percentile of this distribution. We generated a set of randomly directed vectors in PCA space with lengths drawn from a uniform distribution

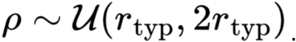

Corresponding hidden-state vectors were obtained by applying the inverse PCA transform. In this way, the sampled states lie on a shell outside the typical wake-state distribution rather than near its center.

We then used Isomap to produce a nonlinear low-dimensional embedding of the wake activity and fitted a nearest-neighbors model on this embedding. Using this model, we calculated distances from each uncharacteristic state to its nearest neighbor in wake activity, and selected 2000 most distant states as initial pRNN states. Starting from each selected initial state, we simulated spontaneous recurrent dynamics for 50 steps, producing uncharacteristic trajectories. For each step in the trajectory, the distance was then calculated using the same nearest-neighbors model.

To assess the spatial representation of networks trained on multiple environments (Figure 5), performance was assessed per-environment by computing the spatial RSA of the hidden-unit activity after simulating a 15,000 step trajectory in only one environment. To visualize the representation (Fig. 5b), a long trajectory was run for each of three environments, and 4000 timepoints were subsampled per environment, and the hidden activity was pooled. This was used to fit an isomap following the same approach as above. 8000 timesteps from each environment were projected onto the embedding and colored by environment identity.

To assess how discriminable the training environments were from network activity, a multinomial logistic regression classifier (scikit-learn LogisticRegression, L-BFGS solver) was trained to predict environment identity from the hidden-unit activation vector at each timestep. Data were split 80/20 into train and test sets (stratified by environment), features were standardized (zero mean, unit variance) using statistics from the training set, and the classifier was evaluated on the held-out test set.

To quantify how the dimensionality of the hidden-unit representation scaled with the number of distinct environments experienced, two complementary measures were computed from the pooled hidden-unit activity of all environments that each net was trained on (1-7, 5 seeds each). For a linear measure, the participation ratio was computed from the eigenvalues of the principal component analysis, computed on the pooled hidden unit activity. A high ratio means all dimensions contribute similarity, while a low ratio implies uneven contribution. The equation is

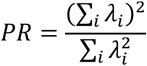

For a non-linear measure, the correlation dimension was estimated using the correlation-integral method (Grassberger & Procaccia, 1983). This is a lower bound on the box-counting dimension (Voigts et al., 2025), but is more tractable, as the compute cost scales linearly with dimensionality (and quadratically with number of data points), while the box-counting dimension cost is exponential with dimension. For a given set of hidden-unit activations, which are 800 dimensional, all pairwise Euclidean distances are computed, and the correlation integral C(r) – the fraction of pairs closer than the radius r – was evaluated at 25 log-spaced radii spanning the 5^th^ – 60^th^ percentile of pairwise distance distribution (chosen to avoide the small-scale noise floor and the large-scale saturation). The correlation dimension is then taken as the slope of the least-squares fit of the log C(r) vs. log r plot.

In Rollout networks, there are k*t activations every time the animal explores an environment for t steps. When metrics such as sRSA and SWdist are computed for rollout networks, the activations at each timestep are averaged over the k rollout steps to arrive at one activation for each timestep of the agent’s exploration.

### Experiments

The following experiments were run on networks after training.

#### Single-trial experience replay

To calculate the sleep reactivation of positions visited during wake experiences, 150 unique wake trials of *T* = 100 timesteps were simulated. The observation sequences from each of these trials were used for a single weight update of a trained network, with learning rates that ranged from 0 to 10^3^ times higher than the learning rate used during training (see Figure S18, learning rate was varied for recurrent weights and biases, only). The outcome of this procedure was 1200 trained networks, which differed only in the data and learning that occurred in the last trial: one of 100 different paths through the environment, at different learning rates. The resulting networks were then each run for 20 adaptation-generated sleep trials of *T_sleep_* = 100 timesteps. The spearman rank correlation between the occupancy of the decoded position, *P*[*x̃_t_*], and the position from corresponding wake trial *P*[***x****_t_*].

#### Object association learning

To test the network’s ability to learn new associations between sensory inputs and positions, the network was placed into a new environment, which was identical to the original with the exception of a single-square bright green [0.3, 0.5, 0.3] “object”. The network was then trained for 200 trials of *T* = 1000 timesteps with learning rate for the output weights doubled. Its ability to remember the new sensory association was assessed by running a *T* = 1000 timestep trial in the original (object-free) environment, and measuring the change of the value of the corresponding green pixel in the network output, ***y****_t_*, when the network was viewing the position where the object had been located.

##### Planning

To test the utility the network’s activity for planning actions towards a goal (**Figure S9**), the following procedure was used:

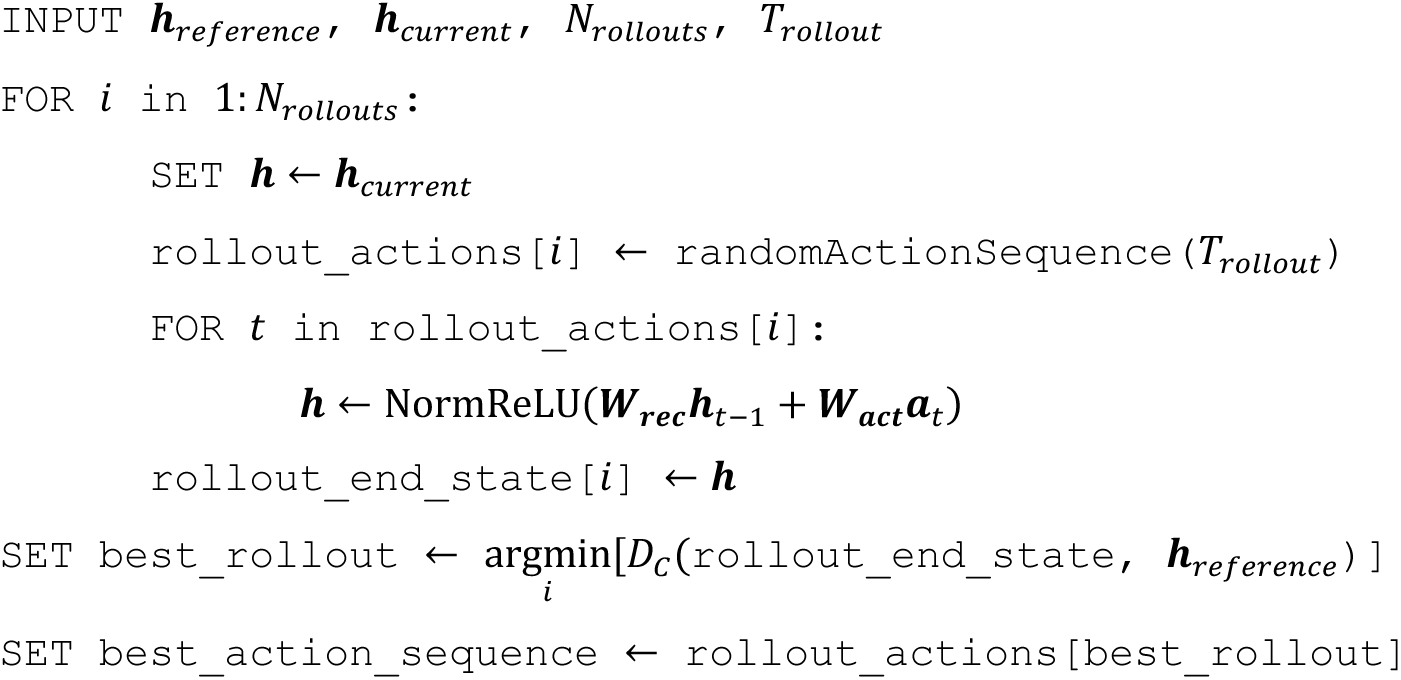

Where ***h****_reference_* is the neural activity corresponding to the goal state, ***h****_current_* is the current neural activity state, and `randomActionSequence(`*T_rollout_*)returns a random action sequence of length *T_rollout_*. The states used as ***h****_reference_* and ***h****_current_* are described in the figure legend for **Figure S10**.

### Environments and agents for testing task-relevant non-spatial representation

To examine the emergence of non-spatial variables, networks were trained as outlined previously in the methods, but with the gridworld environments shown in **Figure 8a,c**, and **g**, and with the following agents, instead of the random action agent.

#### Figure-8 maze alternation agent

At the start of each trial, the agent was initialized at the top of the stem (the middle branch of the figure-8). It then took either a “stop” action, with probability 0.05, or followed the next action in a deterministic policy that caused the agent to follow a figure-8 pattern through the maze. Trials ran for T=600 timesteps

#### Circle maze 4-laps agent

At the start of each trial, the agent was initialized to start at the top left corner of the environment. It then took either a “stop” action, with probability 0.05, or followed the next action in a deterministic policy that cause the agent to make 4 clockwise laps of the environment, ending back at the start position after ∼16 steps (12 forward steps and 4 clockwise turns). Every fourth lap around the environment, the agent took 7 stop actions, during which a “reward” stimulus was presented (see below), followed by 2 more stop actions with no reward stimulus. It then restarted the 4-lap policy for a total of T=600 timesteps. To encode the reward stimulus, the observation at time t corresponded to its 7×7×3 egocentric view as well as a binary pixel which represented the presence or absence of the reward. The reward pixel was broadcasted to 5×7×3 and concatenated to the egocentric view, making it ∼42% of the prediction error signal (147 visual pixels and 105 reward pixels).

This environment was also used to test for a shift in place field center (**Figure S25**). Four networks were first trained for 10,000 trials of the random policy. The deterministic laps policy was then used to measure cell’s spatial tuning curves. Networks were then trained for an additional 10,000 trials with the deterministic policy and spatial tuning curves were again measured. The change in peak location of the place field was compared for cells with at least 50% EV, 0.5 SI, and identified to have a single field both before and after the deterministic training.

#### 3-goal maze tone agent

At the start of each trial, the agent was initialized at the far left of the linear maze, facing right (towards the goals). It then took either a “stop” action, with probability 0.05, or followed the next action in a deterministic policy that cause the agent to proceed in a “short”, “medium”, or “long” lap to the near, middle, or far goal, respectively, and then turn around and return to the start location, at which point it turned around and proceeded with another lap for a total of T=600 timesteps. The sequence of laps was randomized each trial. A “tone” stimulus as added as a single additional pixel to the observation, which increased from 0 to 1 during the forward direction half of the lap, following the agent’s spatial progression to the goal.

## Acknowledgements

The authors would like to thank Antonio Fernandez-Ruiz and Dileep George for helpful comments on the manuscript.

DL: This work was supported by the FRQNT Strategic Clusters Program (2020-RS4-265502 – Centre UNIQUE – Unifying Neuroscience and Artificial Intelligence – Québec) and the Richard and Edith Strauss Postdoctoral Fellowship in Medicine. RHE: Deepmind fellowship. AP&BR: New Frontiers in Research Fund (Exploration grant NFRFE-2021-00926 and HBHL - Healthy Brains for Healthy Lives (Innovative Ideas grant 1c-II-15). BR: This work was supported by a NSERC (Discovery Grant: RGPIN-2020-05105; Discovery Accelerator Supplement: RGPAS-2020-00031; Arthur B. McDonald Fellowship: 566355-2022) and CIFAR (Canada AI Chair; Learning in Machine and Brains Fellowship). This research was enabled in part by support provided by (Calcul Québec) (https://www.calculquebec.ca/en/) and the Digital Research Alliance of Canada (https://alliancecan.ca/en). The authors acknowledge the material support of NVIDIA in the form of computational resources. AP: NSERC (Discovery Grant: RGPIN-2018-04600), CIHR (Project grants 155957 and 180330)

## Author Contributions

DL, AP, BR designed the research. DL, AP, BR wrote the manuscript. DL, AE, RHE carried out analysis.

## Additional Information

Supplementary Information is available for this paper.

Correspondence and requests for materials should be addressed to Blake Richards, Adrien Peyrache, and Dan Levenstein.

## Supplemental Figures

**Figure S1:**
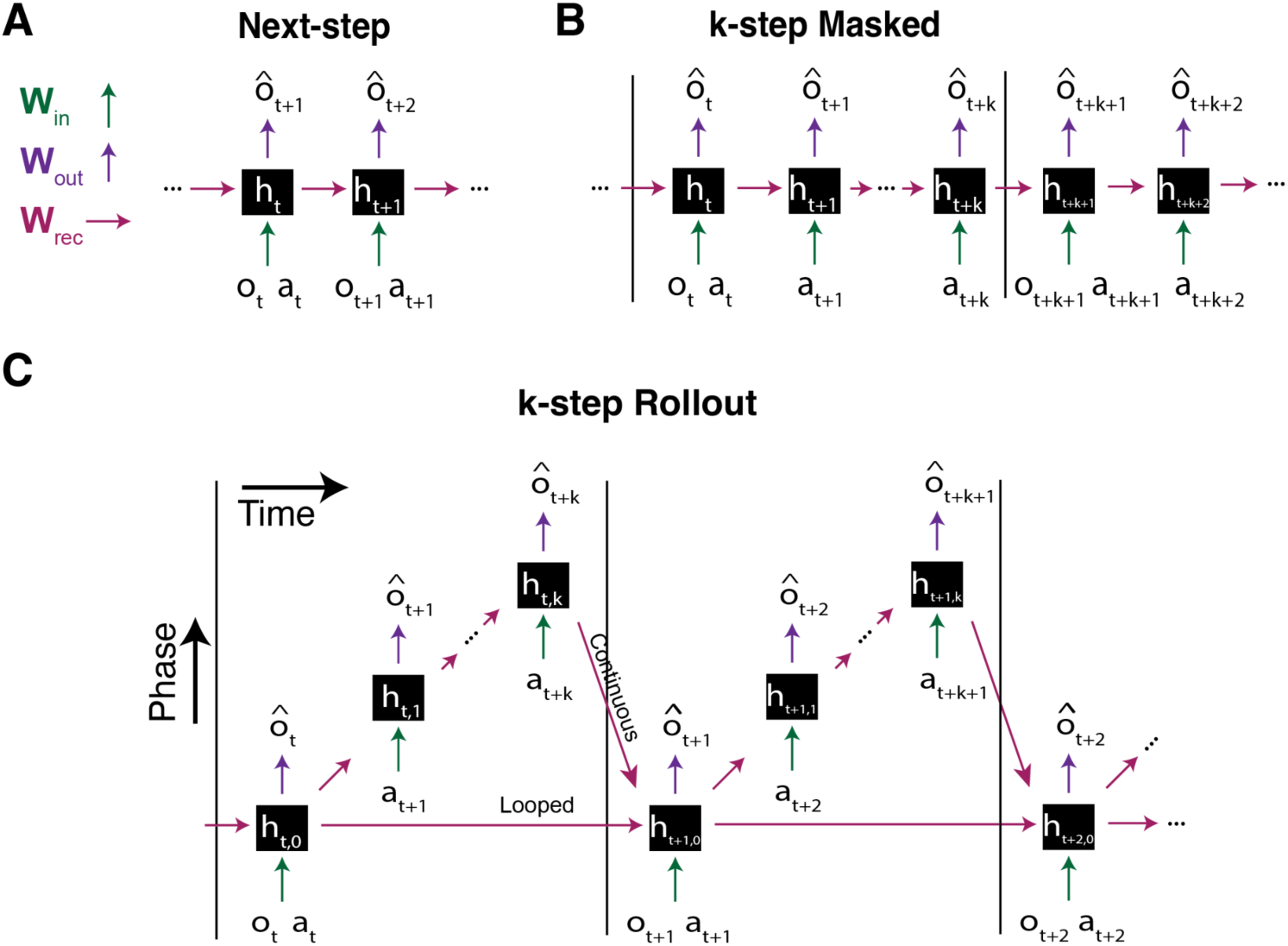
Block diagrams for predictive RNN architectures used in this study. All networks are trained using *T*-length sequences of observations and actions (default *T* = 500), in which *o_t_* is the sensory observation at time *t*, *a_t_* is the subsequent action taken, and *o*i*_t_* is the network’s prediction of *o_t_*. All weights, as well as the bias of the recurrent hidden units, ℎ, are trained using back propagation through time. Further details can be found in the methods. **A:** The next step pRNN is trained to predict the observation at the next time step. **B:** The masked pRNN is trained to predict the observation at the current time step, which is not included in the input except every *k* + 1 time steps. The network is engaged in sequential predictive learning when *k* > 1, as the network must predict *k* time steps of masked inputs in a row. **C:** The rollout pRNN is trained to predict a *k*-length sequence of future observations. Time in the rollout pRNN can be “continuous” (ℎ*_t_*_+1,0_ = *f*(ℎ*_t_*_,*k*_, *o_t_*_+1_, *a_t_*_+1_)) or “looped” (ℎ*_t_*_+1,0_ = *f*(ℎ*_t_*_,0_, *o_t_*_+1_, *a_t_*_+1_)), depending on if the recurrent input in the beginning of the rollout comes from the hidden state at the end or beginning of the previous rollout.

**Figure S2:**
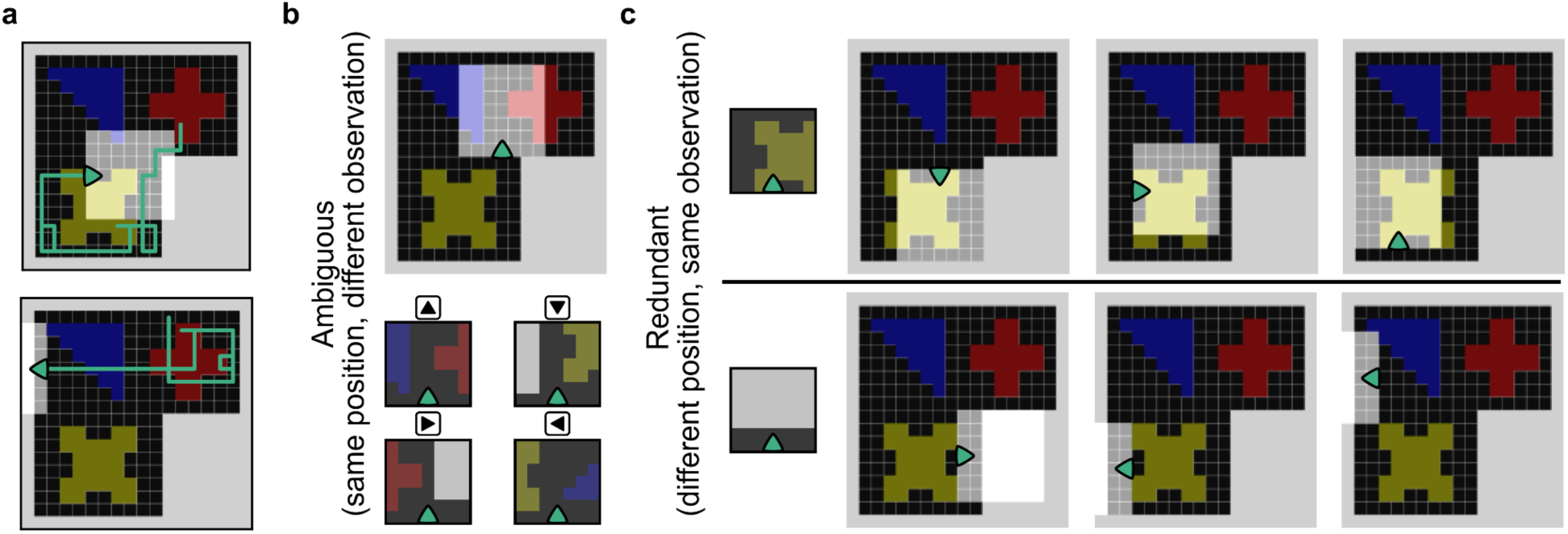
Spatial ambiguity and redundancy of sensory input in the gridworld environment. **a:** Two example 50-step trajectories using the random action policy in the gridworld environment (trajectories used for training are 600 steps unless otherwise specified). **B**: Sensory input in the environment is ambiguous – the same position can have different sensory inputs, depending on the orientation (“head direction”) of the agent. **C**: Sensory input in the environment is redundant – different positions in the environment can have identical sensory inputs.

**Figure S3:**
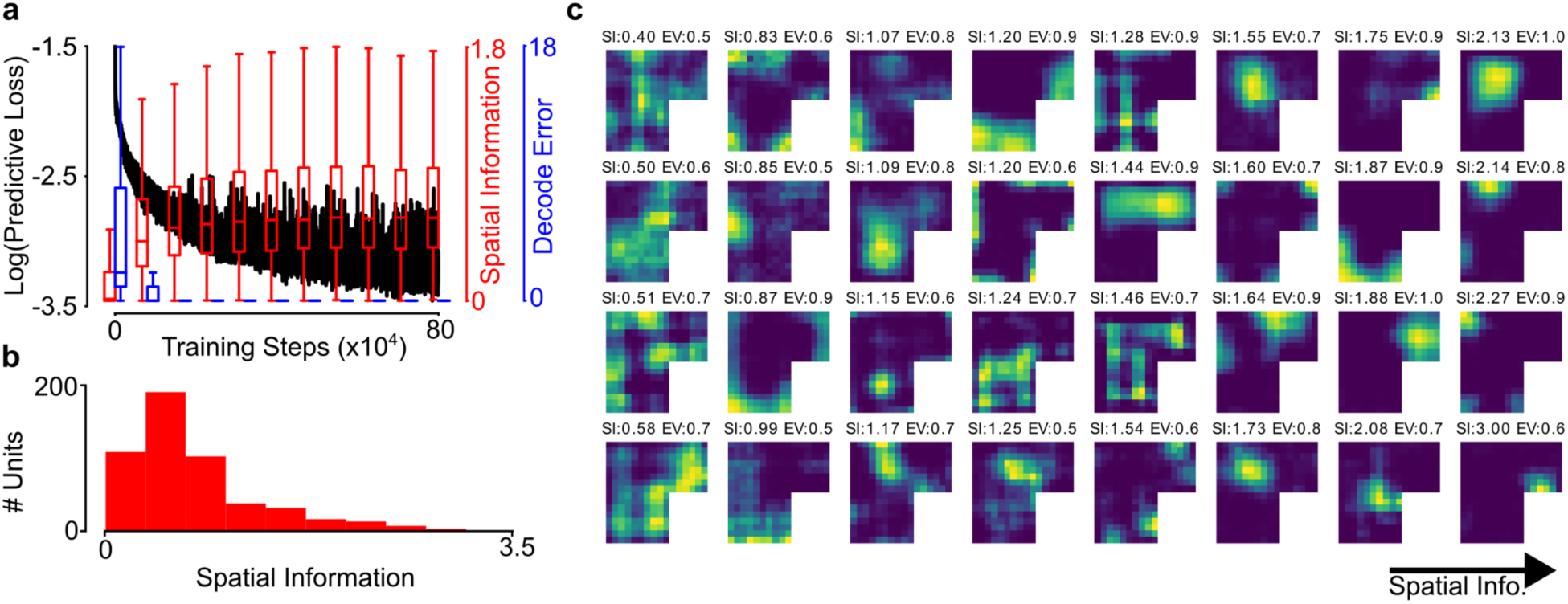
Training and Spatial Tuning in the Next-Step pRNN. **a:** The evolution of predictive loss, distribution of spatial information, and decode error over training. Each training step is a single wake epoch. **B:** Distribution of spatial information over all cells in the trained network. **C:** Tuning curves from a random selection of tuned cells (>50% variance explained by space).

**Figure S4:**
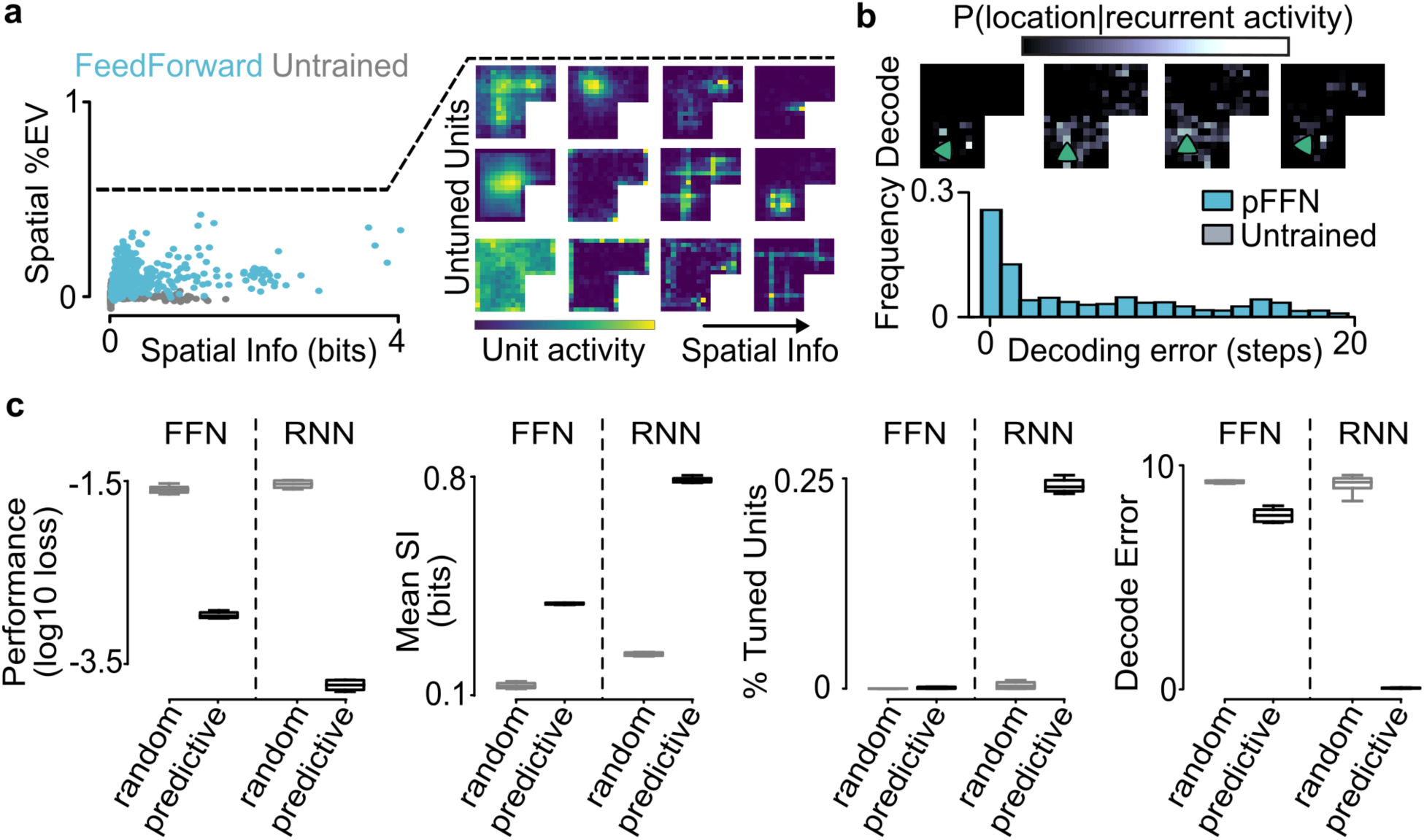
Spatial tuning in feedforward and untrained networks. **a:** (Left) Explained variance by spatial position and spatial information of all units in the example predictive feed-forward network and units from an random feedforward network. (Right) spatial tuning curves from a selection of example units from the network, sorted by the spatial information of their tuning curves. Note that no units have more than 50% of their activity variance explained by the agent’s position. Compare to Figure 2c. **B:** (Top) Decoded position from a linear decoder on hidden unit activity in the feedforward network for four example timesteps. (Bottom) distribution of decoding error over a 1000-timestep trajectory in the environment, for the example feedforward and untrained network. (Compare to Figure 2d) **C:** Predictive performance, mean spatial information, % of tuned cells, and mean decoding error for random and predictive feedforward networks (FFN) and recurrent networks (RNN). Box and whiskers correspond to distribution over 9 seeds.

**Figure S5:**
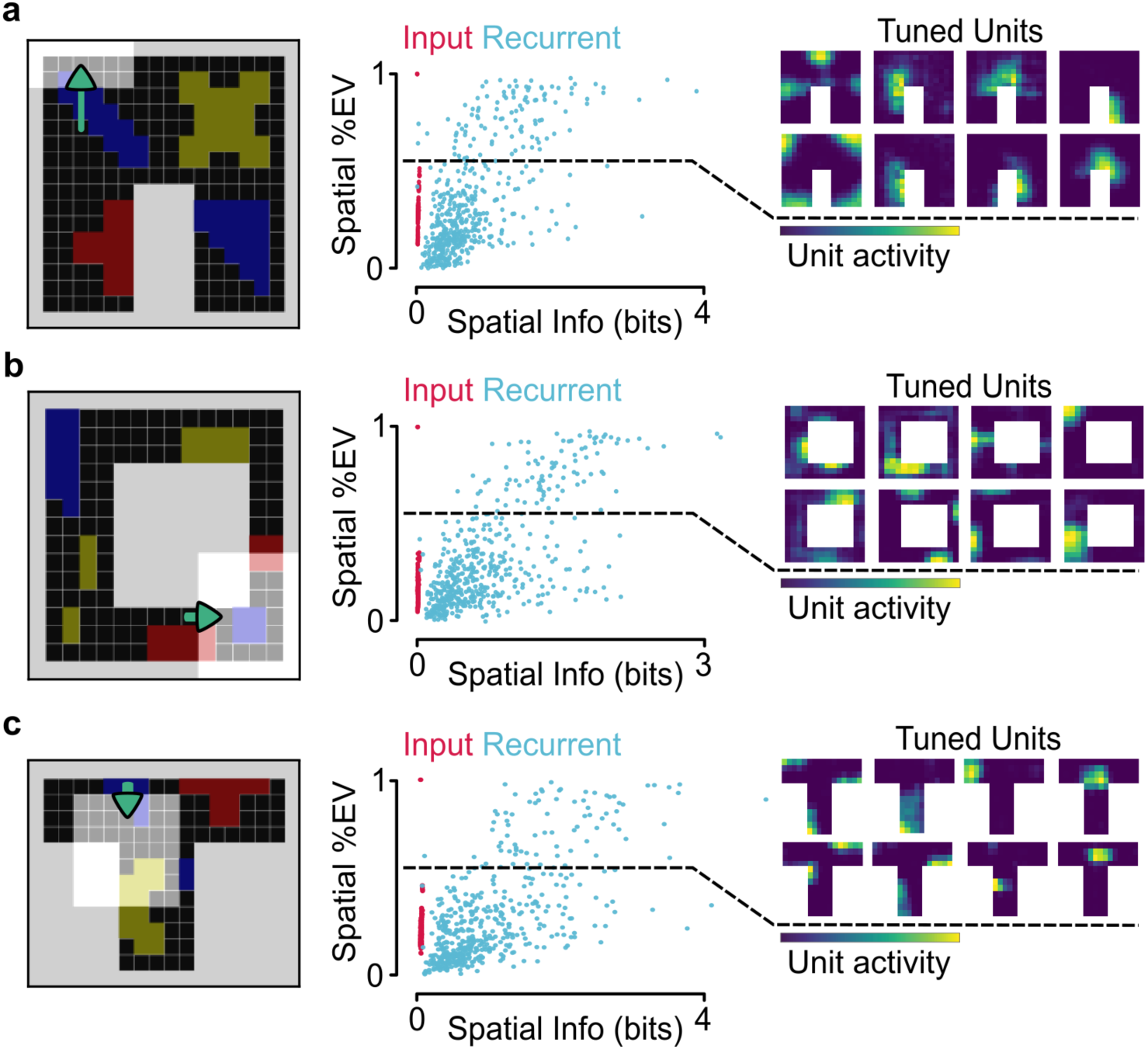
Spatial tuning of recurrent units in gridworld environments with different geometries. **a:** A U-shaped environment. (Left) Top-down view of the environment. (Middle) Variance explained by the agent’s position and spatial information for all recurrent and input units in an example next-step pRNN. (Right) Spatial tuning curves for example tuned units (units with more than 50% variance explained by spatial position). Compare to Figure 2c. **B:** Same as panel a, for a circular environment. **C:** Same as panel a, for a T-shaped environment.

**Figure S6:**
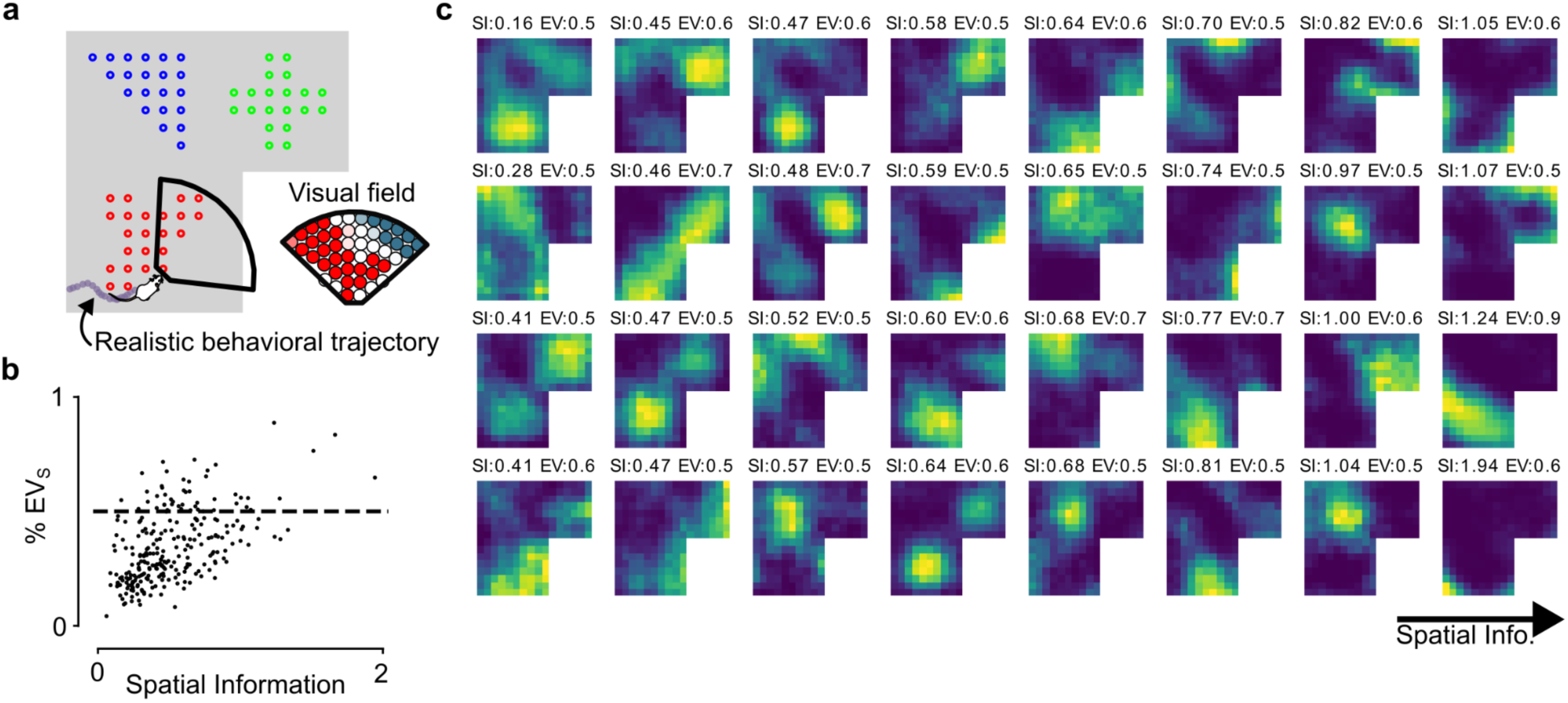
Spatially-tuned cells in a next-step pRNN in the RatInABox environment. **A**: The RatInABoxEnvironment (George et al 2024). An agent takes rodent-like continuous trajectories in the environment, and collects egocentric visual inputs. **B**: %EVs and Spatial information for all units in the network. **C**: Tuning curves from a random selection of tuned cells (>50% variance explained by space).

**Figure S7:**
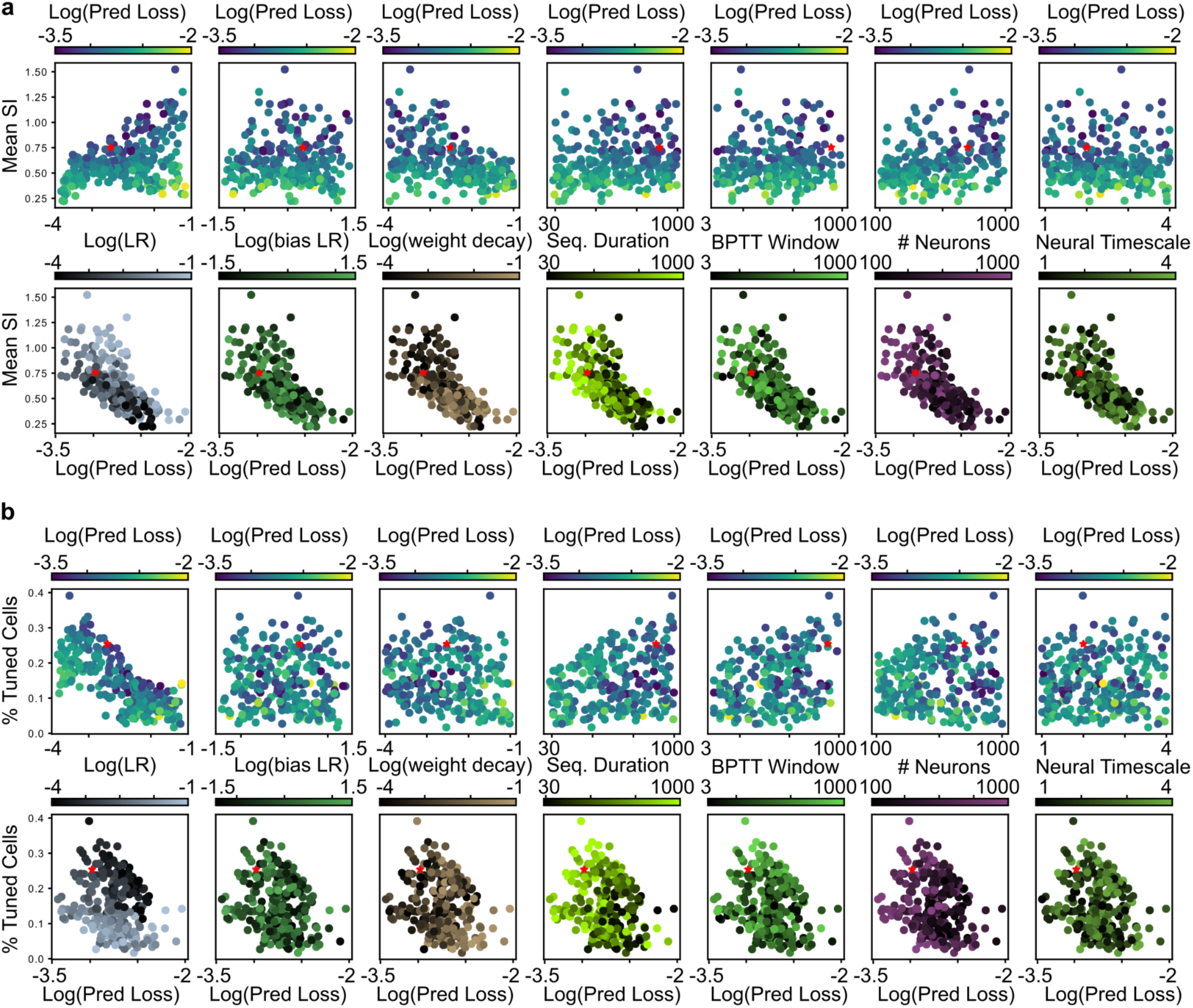
Random hyperparameter population. **a:** (Top) Spatial information and predictive loss (color) for all networks in the random hyperparameter sweep, as a function of each hyperparameter. (Bottom) Spatial information (color) and predictive loss for all networks in the random hyperparameter sweep, as a function of each hyperparameter. **B:** Similar to A, for % Tuned Cells.

**Figure S8:**
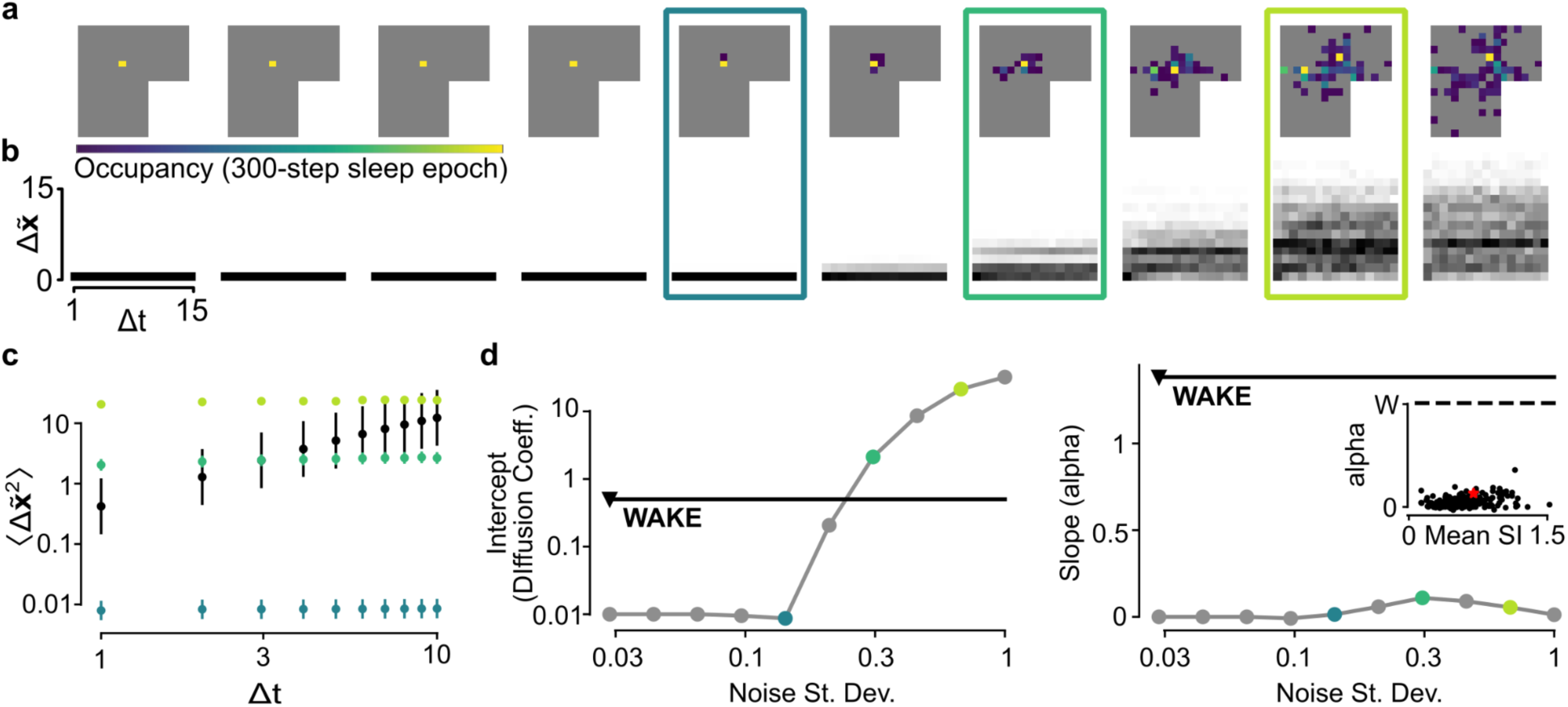
Sleep activity in the Next-step pRNN as a function of the variance of internal noise. **a:** Spatial occupancy histograms for sleep epochs with different levels of noise. Note: each epoch has a different seed for initial hidden unit state and noise generation. **B:** Spatial transition statistics for different levels of noise. **C:** Diffusion plot for three example levels of noise. **D:** Diffusion constant (intercept, right) and alpha (slope, left) of offline activity as a function of noise. Note: due to layer normalization, the standard deviation of recurrent input is 1, black triangle indicates noise level during training, and noise was set to 0.5 for Figure 2 f-i. Inset: alpha for all networks in the random hyperparameter population.

**Figure S9:**
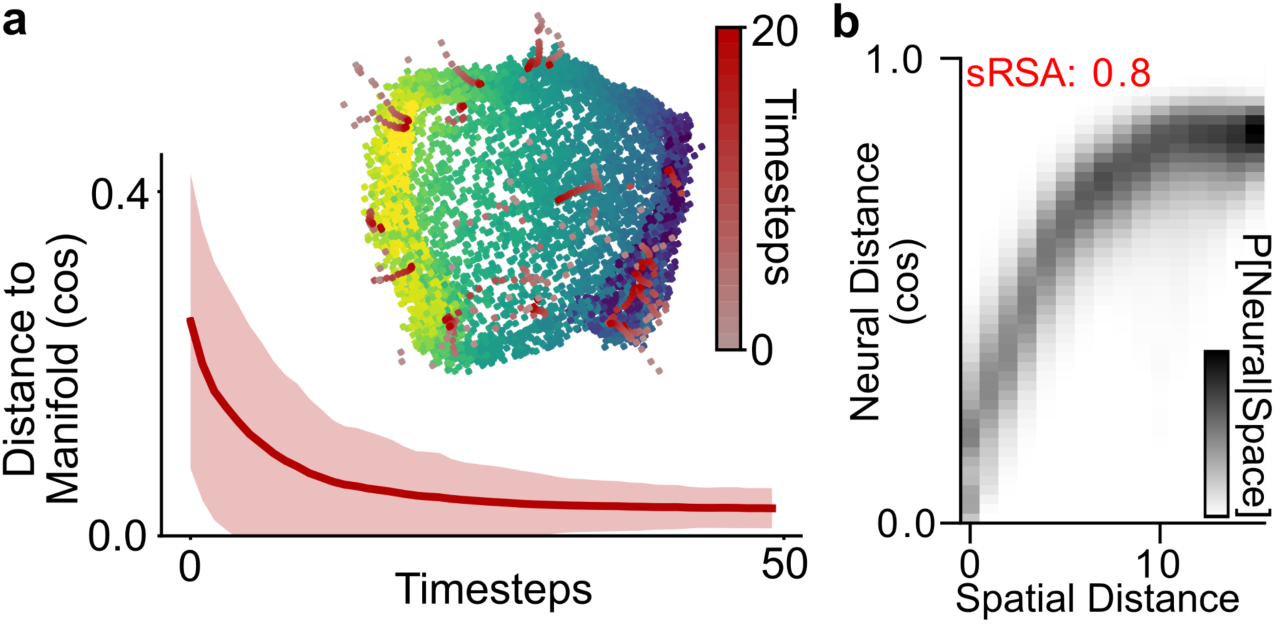
Sequential predictive learning produces a continuous attractor manifold. **a**: Distance to the wake manifold of spontaneous (sleep) trajectories with different random initializations. (Mean +/- std over 100 trajectories). Inset: isomap projection of wake manifold and first 20 timesteps of 15 example trajectories. **b**: Distance between neural activity vectors at different timepoints, conditioned on the spatial distance between the agent at those timepoints.

**Figure S10:**
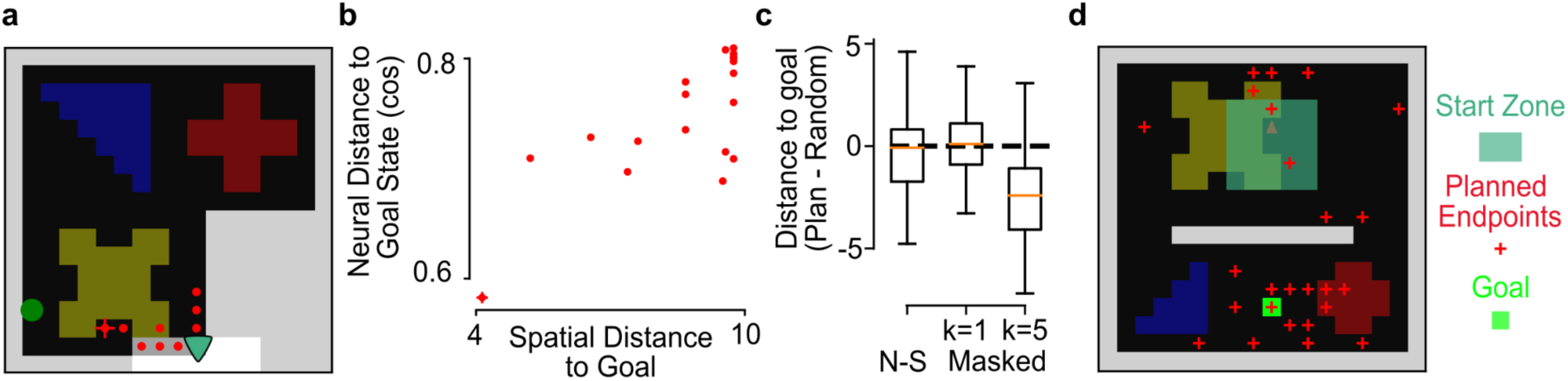
High-sRSA representation of sequential masked networks can act as a cognitive map that informs behavior. **a**: An example session of the navigate-towards-remembered-location task (see Methods). Briefly, the trained network is placed in the environment and allowed to run for a single trial, ending at the green circle; the state of the network is saved. The network is then re-initialized and takes 100 steps to self-localize, ending at the location indicated teal triangle. The network then runs 30 different candidate action plans, each of duration 20 steps, resulting in 30 different network states, and takes the action sequence corresponding to the plan in which the network state is closest in neural space to the remembered goal state (red plus: end of taken action plan, red dots: end of candidate action plans). **b:** Neural distance to goal state at the end of all candidate action plans, as a function of spatial distance to goal position that would result from taking each action plan. **c:** Distance to goal (negative: closer to goal) of 100 different sessions for the next-step, k=1, and k=5 masked networks. Only the k=5 masked network is, on average, closer to the goal as a result of using its internal representation for planning. **d**: Similar to the task in a, but with networks trained in an environment with a wall. The initial state is restricted to the teal box north of the wall, while the goal position is south of the wall. Goal state is compiled from cells’ tuning curves (population vector of average activity at the goal position), rather than a remembered state (see Methods).

**Figure S11:**
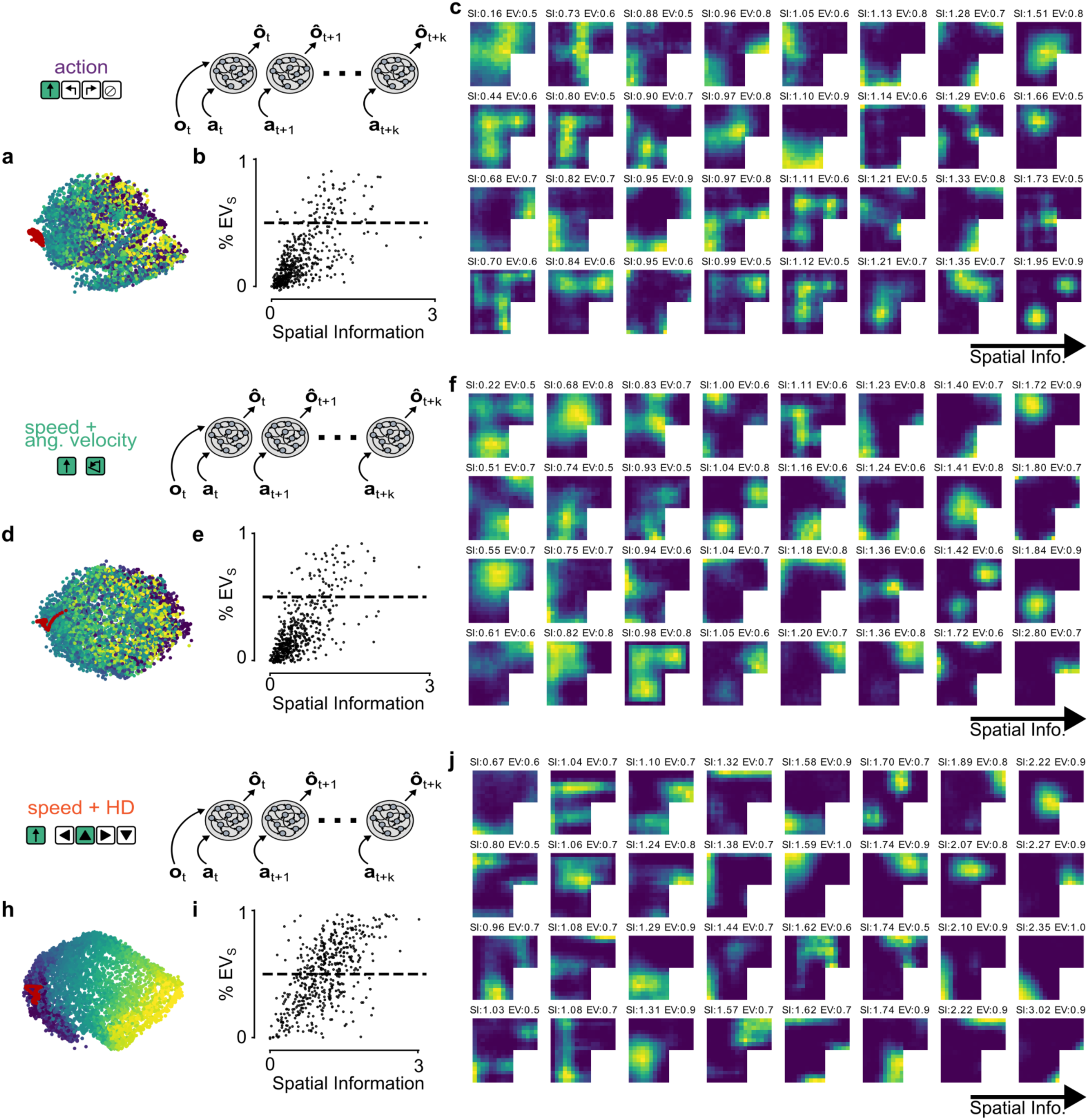
Action-encoding-dependence of cognitive map emergence. **a:** Isomap projection for a 5-step masked network trained with action identity (one-hot encoding). **B:** Spatial information and %EVs for all cells in the network. **C:** Tuning curves from a random selection of tuned cells (>50% variance explained by space). **D-F**: Same as A-C, for a 5-step masked network trained with speed and angular velocity. **H-J:** Same as A-C, for a 5-step masked network trained with speed and head direction (different seed from figure 3C,D).

**Figure S12:**
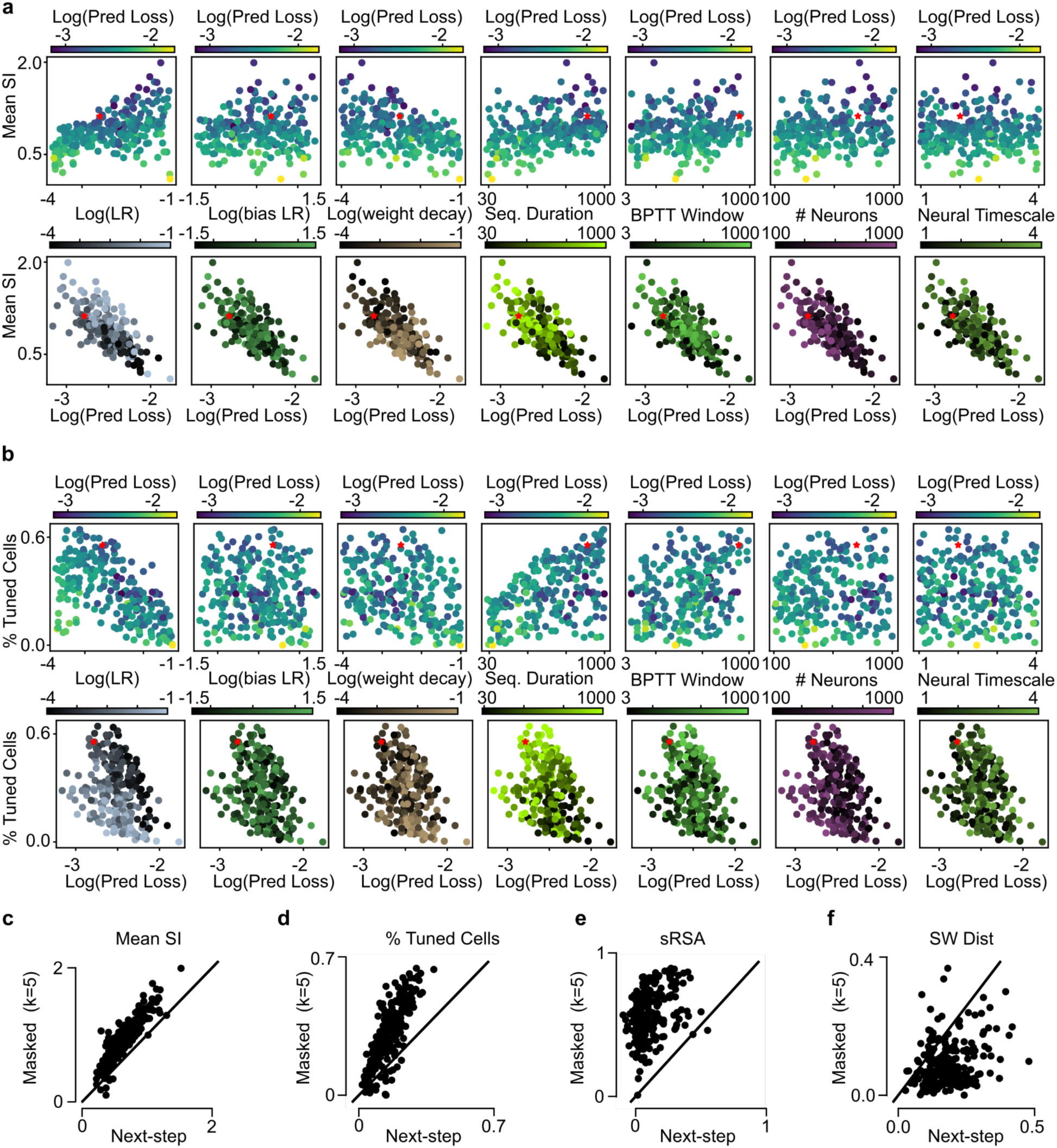
Random hyperparameter population - 5-step masked networks. **a:** (Top) Spatial information and predictive loss (color) for all networks in the random hyperparameter sweep, as a function of each hyperparameter. (Bottom) Spatial information (color) and predictive loss for all networks in the random hyperparameter sweep, as a function of each hyperparameter. Compare to S7a **B:** Similar to A, for % Tuned Cells. Compare to S7b. **C:** Comparison of mean spatial information between 5-step masked and next-step networks with otherwise identical hyperparameters. **D:** Same as c, for % tuned cells. **E:** Same as c, for sRSA. **DF:** Same as c, for SW Dist.

**Figure S13:**
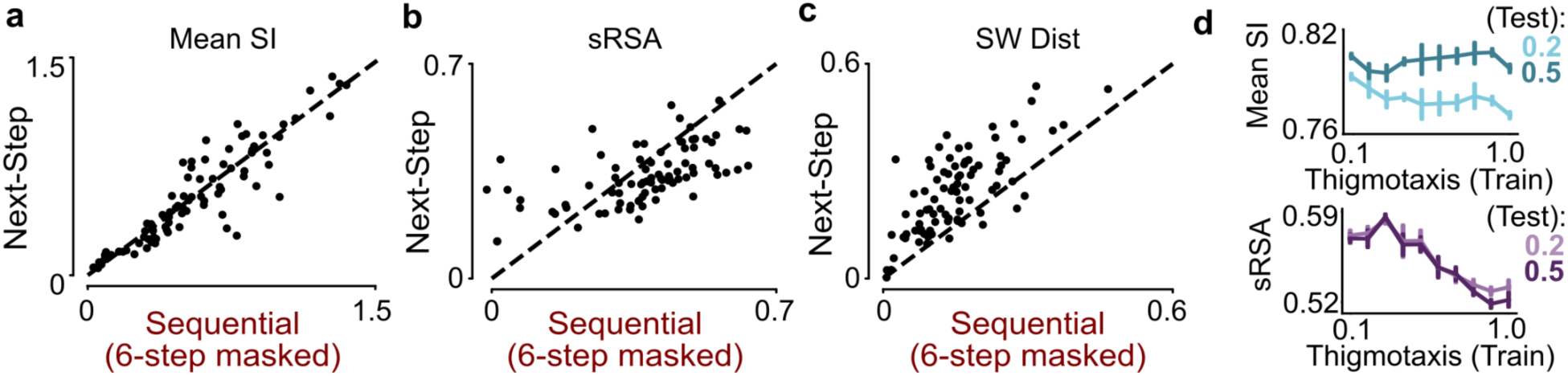
Rat in a box - Sequential vs Next step. **a:** Comparison of mean spatial information between 5-step masked and next-step networks trained in the RatInABox environment, with otherwise identical hyperparameters. **B:** Same as a, for % tuned cells. **C:** Same as a, for sRSA. **D:** mean SI (left) and sRSA (right) of 5-step masked pRNNs as a function of the thigmotaxis of behavioral trajectories used to train the network and during testing.

**Figure S14:**
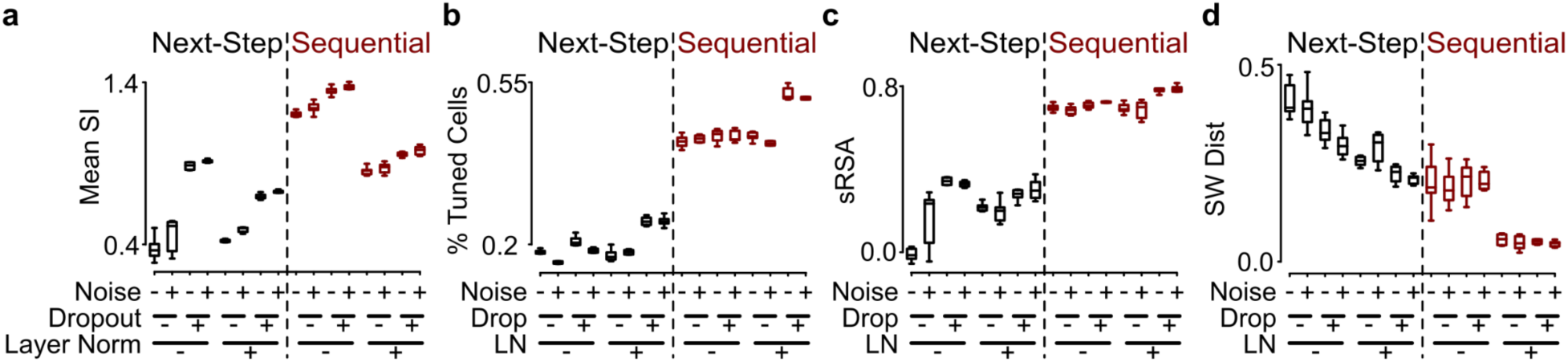
The effect of layer normalization, noise injection, and dropout A-D: A comparison of mean spatial information (**a**), % of tuned cells (cells with >50% variance explained by spatial position, **b**), sRSA (**c**), and SW dist (**d**), across 9 different seeds of next-step and 5-step masked pRNNs trained with/without noise injection, dropout, and layer normalization (see Methods).

**Figure S15:**
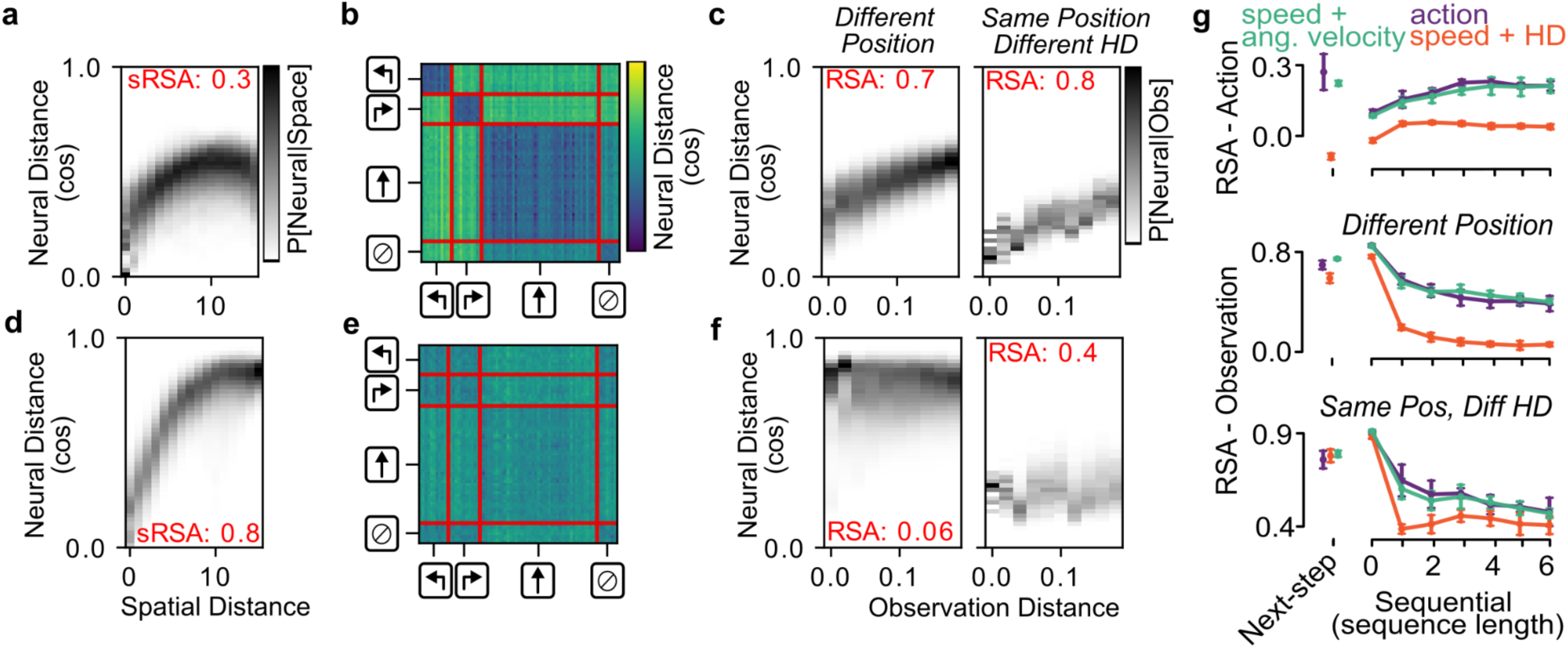
Neural representation of action, observation, and position in next-step and 5-step masked pRNN. **a:** Distribution of neural distance between hidden unit activations at different timepoints, conditioned on the spatial distance between the agent’s position at those timepoints. **B:** Neural distance between hidden unit activations of a next-step pRNN at different timepoints, sorted by action identity. **C:** Distribution of neural distance between hidden unit activations at different timepoints, conditioned on the (pixel) distance between the observations at those timepoints, for distant positions (left), and for the same position but different head direction (right). **D-F:** Same as A-C, for a 5-step masked pRNN. **G:** Representational similarity analysis (RSA) of the relationship between neural activity in pRNNs and action identity (top), observation in different positions (>4 steps apart, middle), and observation in the same position but different head direction (bottom). For action, RSA is the correlation between the neural distance matrix (b,e), and a reference matrix in which same-action pairs are of distance=0 and different-action pairs are of distance=1. For observation, RSA is the correlation between distance in neural space and distance in observations space (c,f). pRNNs include next-step pRNNs and masked pRNNs with k=0 (autoencoder), k=1 (single-step), and k=2-6 (sequential), each trained with one hot (action), speed+HD, and speed+AV encoding of the action input.

**Figure S16:**
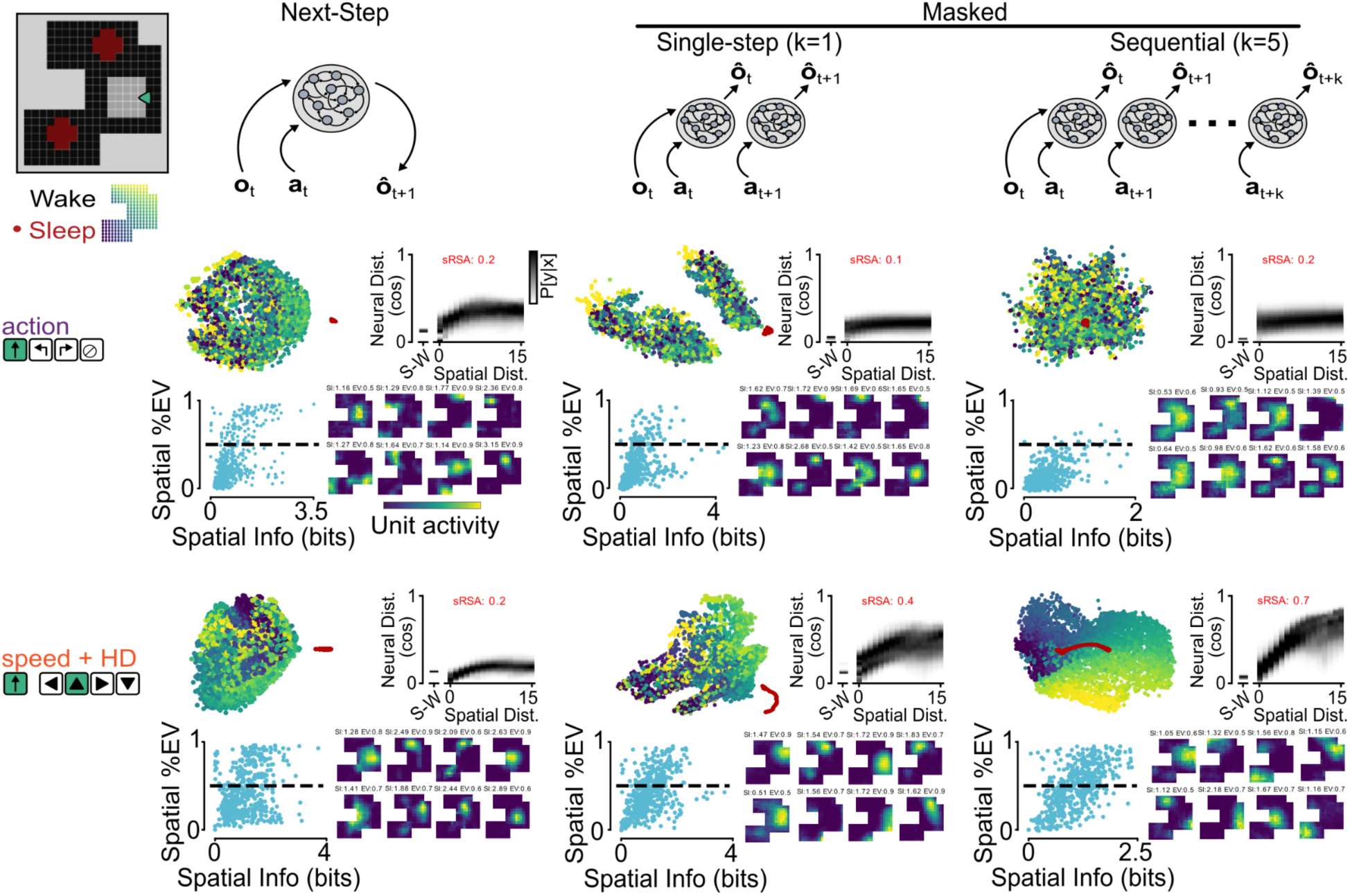
Spatial tuning and cognitive map in an environment with more sensory redundancy. Spatial representation and neural manifold in a gridworld environment with a high degree of redundancy (aliasing). In this environment, the visual field is reduced to a 5×5 grid in front of the agent, and many more positions have identical observation due to the identical floor cues and unmarked floor locations. Comparison is made between next-step (left), single-step masked (k=1, middle), and sequential masked (k=5, right) pRNNs, each trained with action input encoded as action identity (top) or speed+HD (bottom).

**Figure S17:**
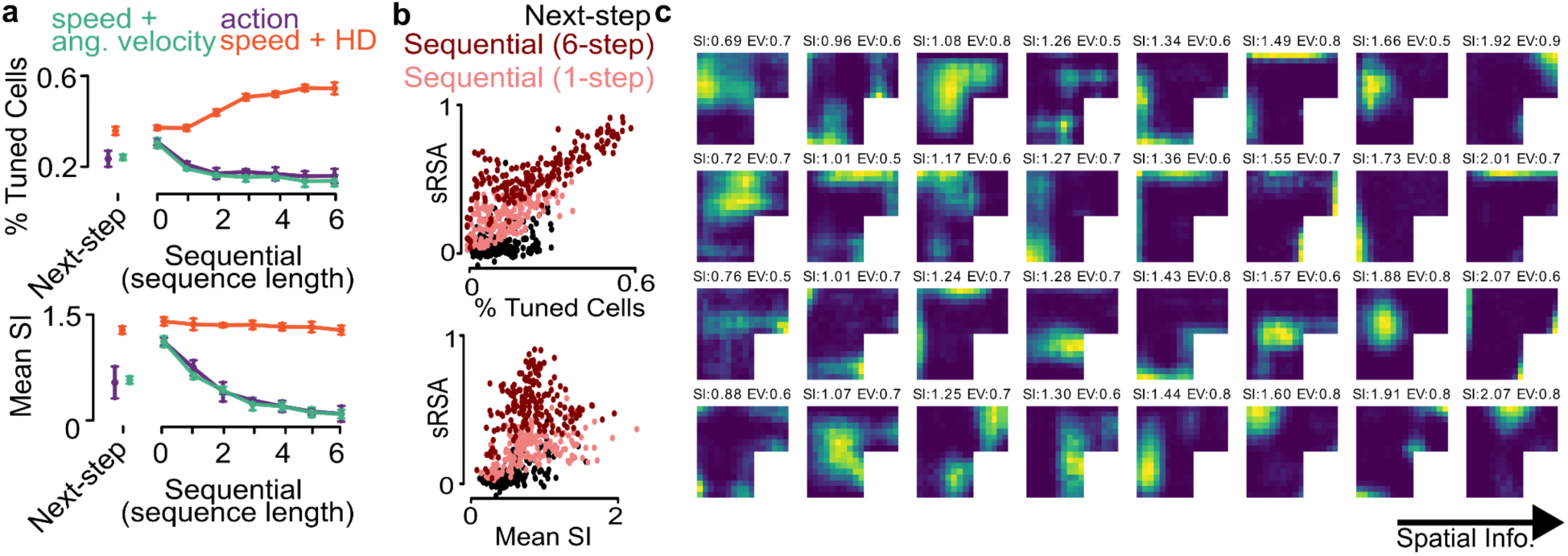
Percentage of spatially tuned cells relates to cognitive map formation. **A**: % Tuned cells (top) and mean SI (bottom) for next-step pRNNs and masked pRNNs with k=0 (autoencoder), k=1 (single-step), and k=2-6 (sequential), each trained with one hot (action), speed+HD, and speed+AV encoding of the action input, at. Mean and standard deviation calculated over 9 seeds. **B:** sRSA as a function of % Tuned cells (top) and mean spatial information (bottom) for the random hyperparameter population of Next-step, single-step, and 5-step masked pRNNs. **C**: Tuning curves from a random selection of tuned cells (cells >50% variance explained by space).

**Figure S18:**
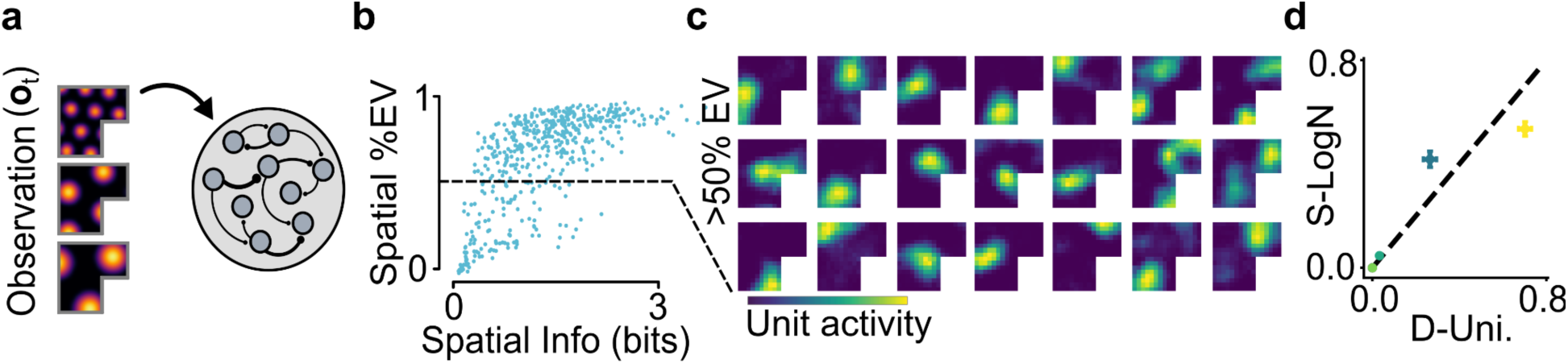
Grid cell input. **a:** A version of the network, using the RatInABox package, in which the observation (input and predicted) corresponds to simulated grid cells rather than the egocentric visual information from the environment (see Methods). **b**: Similar to **Figure 4d**, for the grid cell observation network. **c:** similar to **4b**, for the grid cell observation network. **d**: Similar to **4g**, for the grid cell observation network.

**Figure S19:**
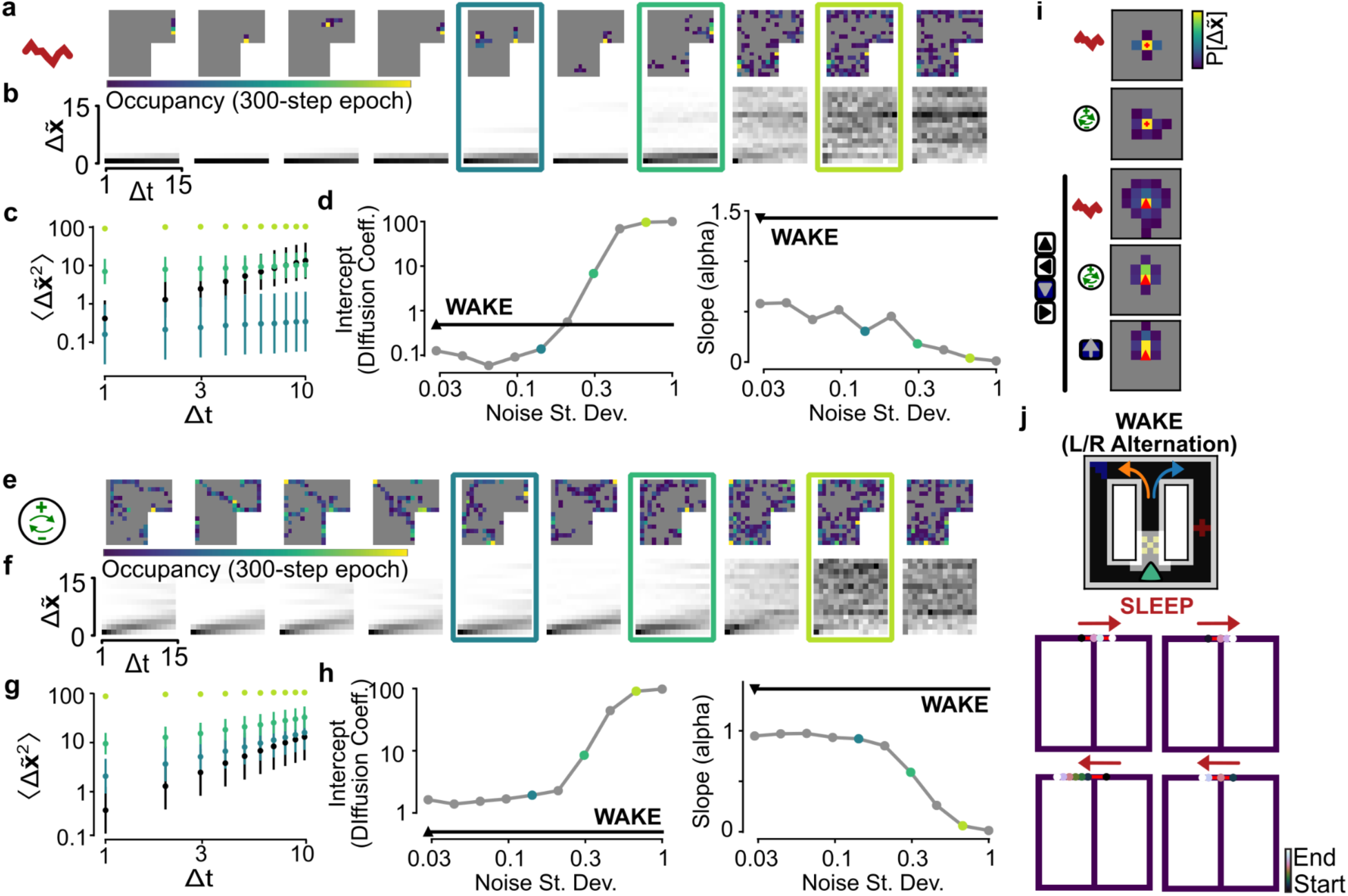
Sleep activity in the 5-step masked pRNN as a function of the variance of internal noise. A-D: Similar to Figure S8 A-D, for a 5-step masked pRNN. **A:** Spatial occupancy histograms for sleep epochs with different levels of noise. Note: each epoch has a different seed for initial hidden unit state and noise generation. **B:** Spatial transition statistics for different levels of noise. **C:** Diffusion plot for three example levels of noise. **D:** Diffusion constant (intercept, right) and alpha (slope, left) of offline activity as a function of noise. Note: due to layer normalization, the standard deviation of recurrent input is 1, black triangle indicates noise level during training, and noise was set to 0.2 for all +noise conditions in **Figure 6**. **E-H:** Similar to A-D, but with adaptation-generated sleep activity. **I:** 1-step transition probability for decoded position during noise-generated, adaptation-generated, and query-directed sleep activity. **J:** Decoded trajectories during sleep that take unseen paths across the top of a maze in which the agent was trained to alternate left and right turns.

**Figure S20:**
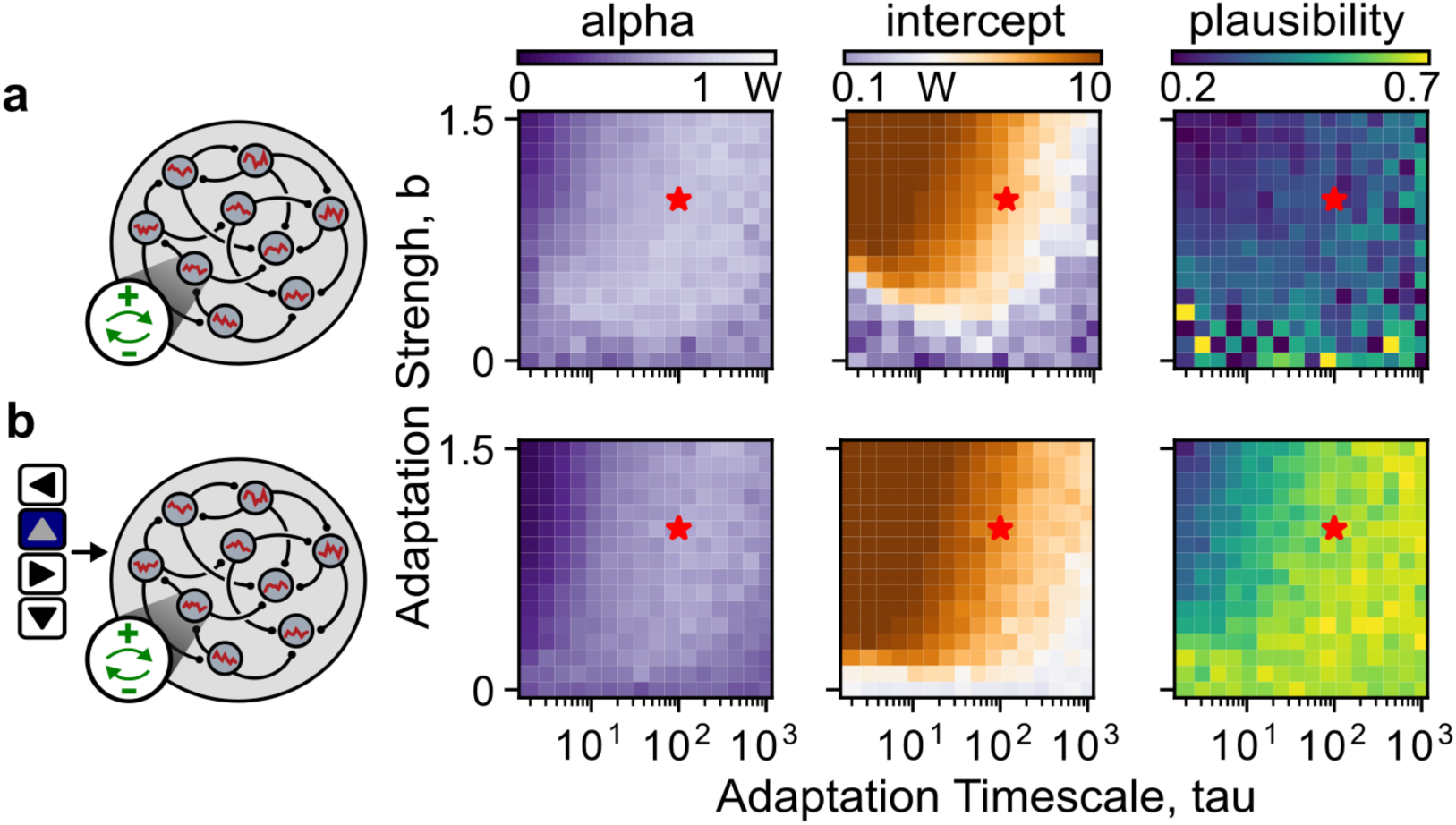
The effect of adaptation parameters on offline trajectory statistics and plausibility. **a:** Alpha (left), intercept (middle), and output plausibility (right) of offline activity as a function of the adaptation strength and timescale during adaptation-driven offline activity in the 5-step masked pRNN. **B:** Same as A, with the head direction query.

**Figure S21:**
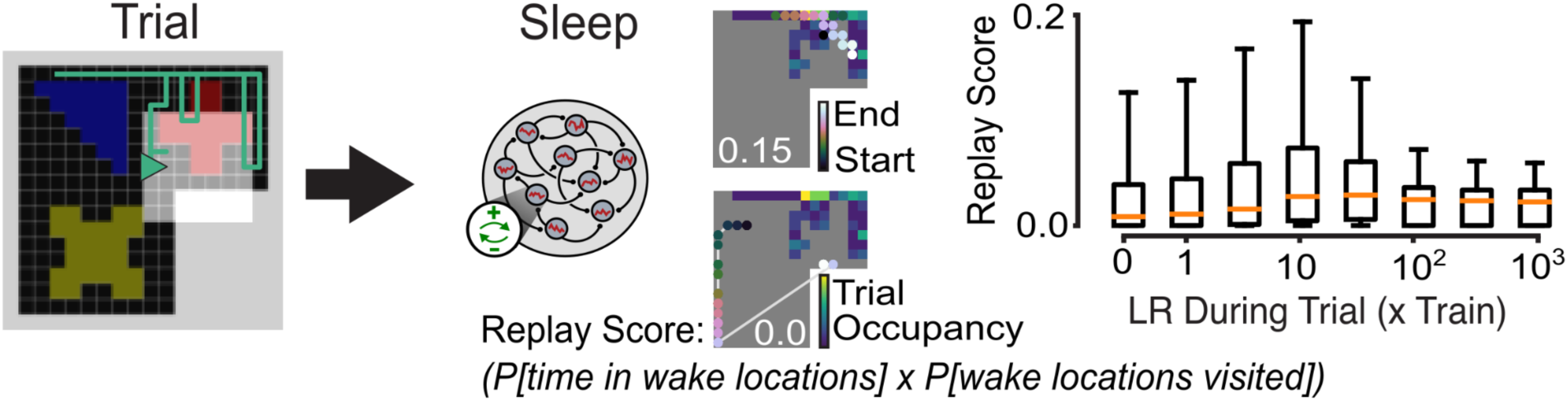
Reactivation of positions from a single-trial. **a:** The trained agent is trained for a single additional wake trial of 100 timesteps. It is then run for a single 100-timestep trial of adaptation-driven sleep activity. Replay score is calculated as (the fraction of time in the sleep trial spent in locations from the wake trail) x (the fraction of wake locations visited). Therefore, a sleep trial in which the sleep visits exactly the same positions as the agent was in during wake will have a replay score of 1. Two example sleep trials are shown for the example wake trial, one with a high replay score and one with a replay score of 0. **B:** Distribution of replay scores over 3000 sleep trials as a function of the learning rate (recurrent weights and biases only) during the wake trial.

**Figure S22:**
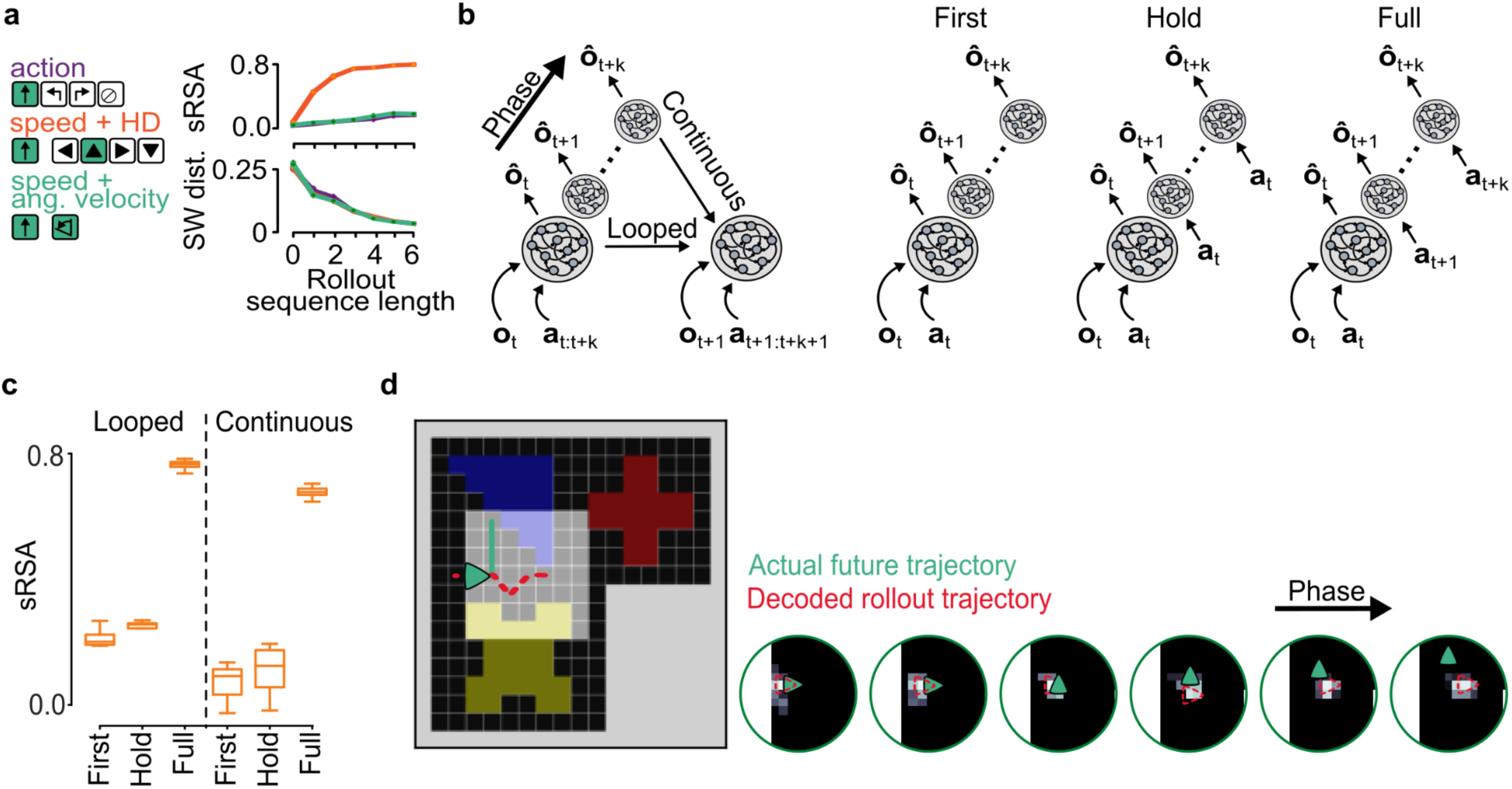
Rollout-based pRNN needs to be trained with true future actions, but once trained can use untaken action to simulate possible trajectories. **a:** sRSA and SW Dist for rollout pRNNs with k=0 (autoencoder), k=1 (single-step), and k=2-6 (sequential), each trained with one hot (action), speed+HD, and speed+AV encoding of the action input, **at**. Mean and standard deviation calculated over 9 seeds. Similar to **Figure 3E**. **B:** Rollout-based pRNN can be trained either with looped or continuous time, and with actions only presented at the first timestep of the rollout (first), with the first action throughout the rollout (hold), or with the actual future action sequence (full). **C:** Comparison of sRSA in networks trained with looped/continuous time and first/hold/full actions provided during the rollout. Only networks trained with the full action sequence develop a cognitive map, per high sRSA, but cognitive map emerges whether time is looped or continuous. Box/whisker indicates distribution over 6 seeds. **D:** An example 5-step rollout in a trained pRNN. Action sequence provided to the network indicated forward movement for the entire rollout, while the actual future actions involved a left turn (teal). Decoded network activity during the rollout continues ahead of the agent (red), diverging from the actual future trajectory.

**FIgure S23:**
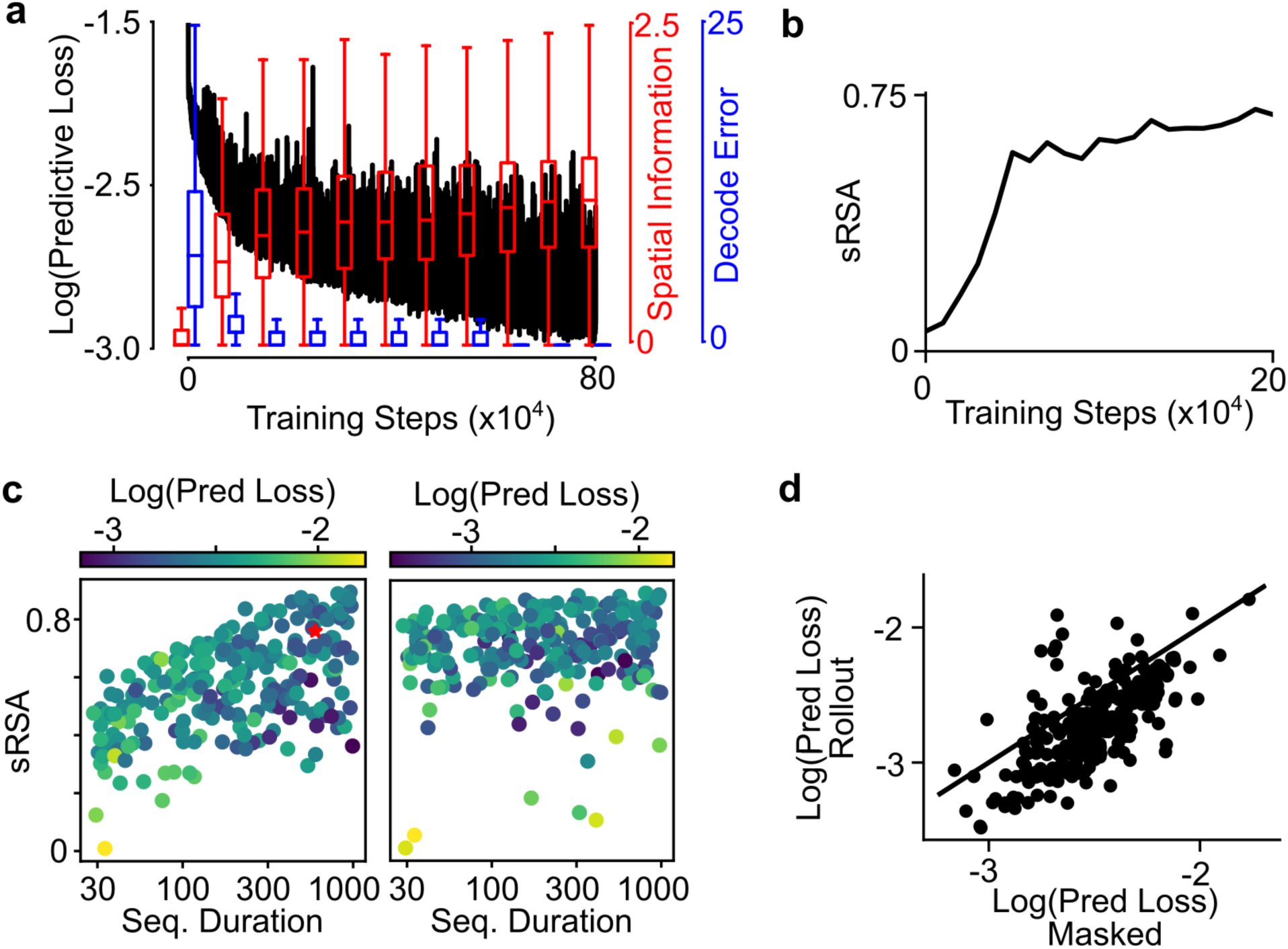
Rollout-based sequential pRNN learns higher-sRSA representation with shorter sequences, and better sensory prediction. **a:** Same as S1A, for a 5-step masked network. **B:** Evolution of sRSA over the first 20,000 timesteps of training for the network in A. Note that sRSA reaches a gradual plateau at ∼5,000 timesteps. **C:** sRSA for all networks in the random hyperparameter population of 5-step masked (left) and 5-step rollout (right) networks, as a function of the duration of sequences used for training. **D:** Predictive loss for hyperparmeter-matched 5-step masked and rollout-based sequential pRNNs.

**FIgure S24:**
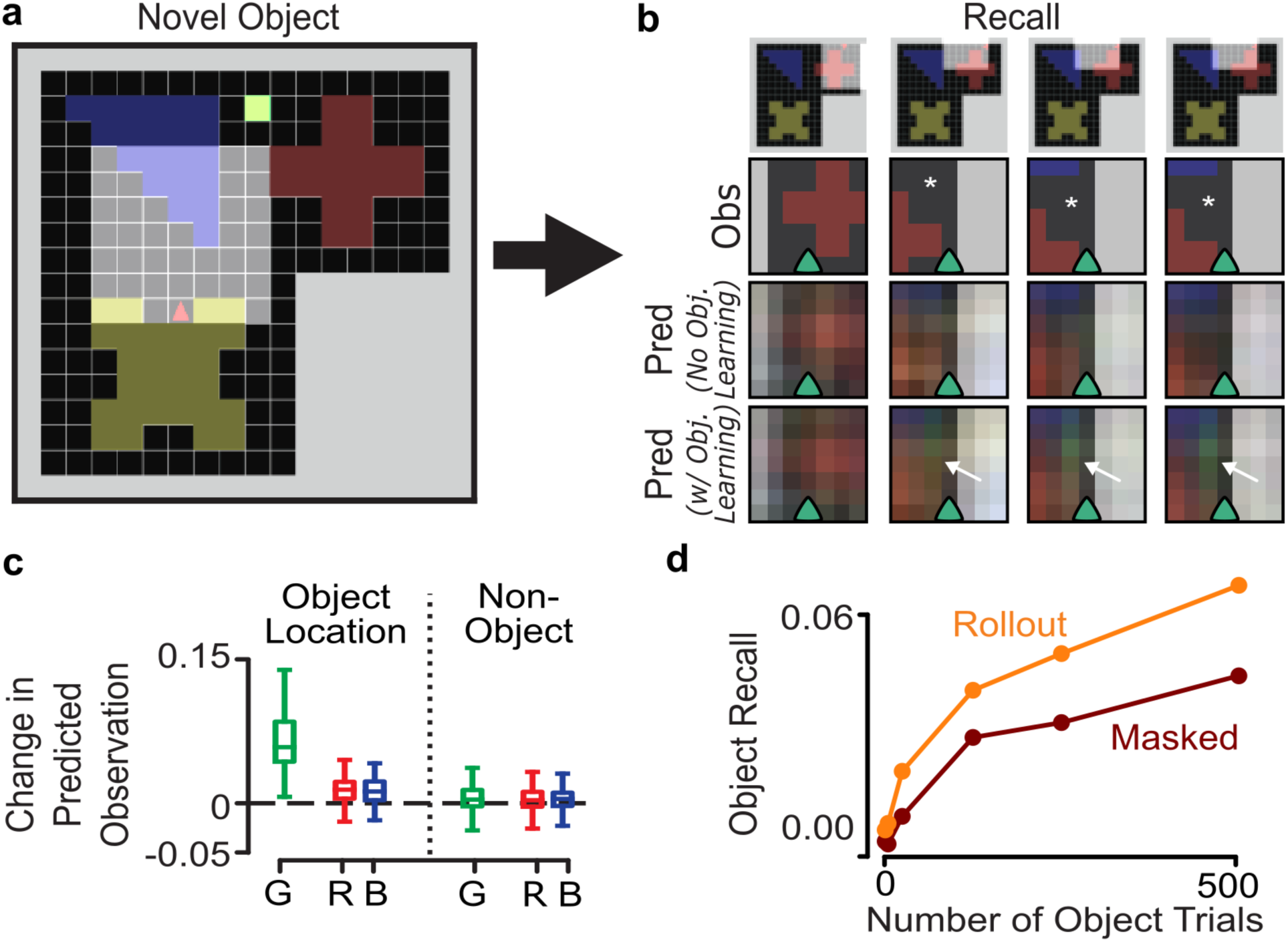
Rollout-based sequential pRNN can quickly learn and recall novel stimuli. A: The novel object environment. After training, the network is exposed to an environment with a novel object (green square), and trained for a small number of wake epochs. B: After novel object training, the agent is placed back in the original environment (no object), and the activation in the green pixel at the object’s location (white star) is compared to the network before object exposure. C: Difference between output of object-trained and original network, for each pixel color at the object location and non-object locations. D: Comparison of object recall (change in green pixel value at object location) between rollout-based and masked sequential (k=5) predictive networks, after a given number of trials in the novel object environment.

**FIgure S25:**
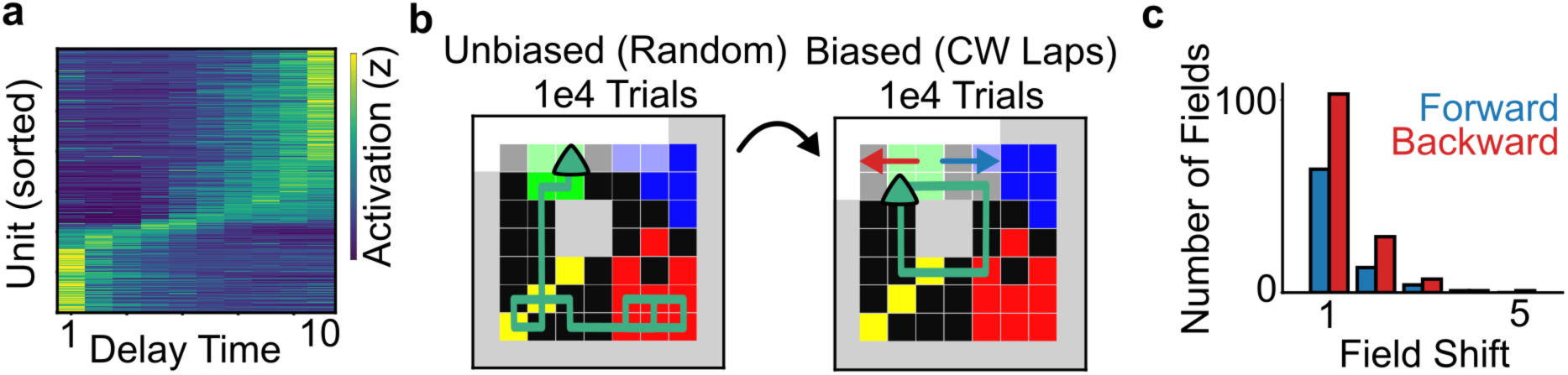
Time Cells appear during the waiting period in the multi-lap environment. **a:** Tuning curve for units in the network with more than 0.5bits/sec about the time interval, as a function of time during the goal-period delay. Units are sorted by the time of their peak activation. Note cells with a peak of increased activity at different times that span the delay period. **B:** The network is first trained with unbiased (random) behavior, followed by training with biased behavior in which it takes clockwise laps around the track. **C**: Direction of place field movement after biased behavior training. Place fields are more likely to shift backwards than forwards.

